# Collective Mechanical Responses of Cadherin-Based Adhesive Junctions as Predicted by Simulations

**DOI:** 10.1101/2021.07.29.454068

**Authors:** Brandon L. Neel, Collin R. Nisler, Sanket Walujkar, Raul Araya-Secchi, Marcos Sotomayor

**Affiliations:** Department of Chemistry and Biochemistry, The Ohio State University, 484 W 12^th^ Avenue, Columbus, OH 43210; The Ohio State Biochemistry Program, The Ohio State University, 484 W 12^th^ Avenue, Columbus, OH 43210; Biophysics Graduate Program, The Ohio State University, 484 W 12^th^ Avenue, Columbus, OH 43210; Chemical Physics Graduate Program, The Ohio State University, 484 W 12^th^ Avenue, Columbus, OH 43210; Facultad de Ingenieria y Tecnologia, Universidad San Sebastian, Chile

## Abstract

Cadherin-based adherens junctions and desmosomes help stabilize cell-cell contacts with additional function in mechano-signaling, while clustered protocadherin junctions are responsible for directing neuronal circuits assembly. Structural models for adherens junctions formed by epithelial cadherin (CDH1) proteins indicate that their long, curved ectodomains arrange to form a periodic, two-dimensional lattice stabilized by tip-to-tip *trans* interactions (across junction) and lateral *cis* contacts. Less is known about the exact architecture of desmosomes, but desmoglein (DSG) and desmocollin (DSC) cadherin proteins are also thought to form ordered junctions. In contrast, clustered protocadherin (PCDH) based cell-cell contacts in neuronal tissues are thought to be responsible for self-recognition and avoidance, and structural models for clustered PCDH junctions show a linear arrangement in which their long and straight ectodomains form antiparallel overlapped *trans* complexes. Here we report all-atom molecular dynamics simulations testing the mechanics of minimalistic adhesive junctions formed by CDH1, DSG2 coupled to DSC1, and PCDHγB4, with systems encompassing up to 3.7 million atoms. Simulations generally predict a favored shearing pathway for the adherens junction model and a two-phased elastic response to tensile forces for the adhesive adherens junction and the desmosome models. Complexes within these junctions first unbend at low tensile force and then become stiff to unbind without unfolding. However, *cis* interactions in both the CDH1 and DSG2-DSC1 systems dictate varied mechanical responses of individual dimers within the junctions. Conversely, the clustered protocadherin PCDHγB4 junction lacks a distinct two-phased elastic response. Instead, applied tensile force strains *trans* interactions directly as there is little unbending of monomers within the junction. Transient intermediates, influenced by new *cis* interactions, are observed after the main rupture event. We suggest that these collective, complex mechanical responses mediated by *cis* contacts facilitate distinct functions in robust cell-cell adhesion for classical cadherins and in self-avoidance signaling for clustered PCDHs.

**Statement of Significance:** Proteins that mediate cell-cell contacts often form aggregates *in vivo* where the tight packing of monomers into junctions is relevant to their function. Members of the cadherin superfamily of glycoproteins form large complexes in which their long ectodomains interact to mediate cell-cell adhesion. Here, we employ simulations to elucidate complex mechanical responses of five junction systems in response to force. Our results offer atomistic insights into the behavior of these proteins in a crowded physiological context, suggesting that classical cadherin complexes in adherens junctions and desmosomes act as molecular shock absorbers with responses modulated by dynamic lateral contacts, while clustered protocadherins form brittle junctions that upon stretching and unbinding form transient interfaces suitable for their critical role in neuronal self-recognition.

## INTRODUCTION

Epithelial and cardiac tissues are subject to perpetual stress from routine physiological stretching and shearing as well as to external forces from cuts and abrasions. To maintain tissue integrity in the face of such relentless perturbations and to sense and respond to mechanical forces, the cells that comprise these tissues have developed strong and mechanically robust cadherin-mediated contacts with one another, such as those formed by adherens junctions and desmosomes (Fig. 1 *A*) (1, 2, 11–20, 3–10). The adherens junction, apical to the desmosome and found throughout the animal kingdom, is formed by homodimerization of epithelial cadherin (CDH1) proteins from opposing cells (Fig. 1 *B*) (1, 21). Desmosomes, universal to vertebrate tissues, are formed by the heterophilic and homophilic dimerization of monomers from two cadherin subfamilies that include desmoglein (DSG) and desmocollin (DSC) proteins (Fig. 1 *E*) (3, 22–26). Failure of CDH1, DSG, and DSC proteins to form native contacts, caused by genetic mutations or autoimmune disease, is associated with a variety of cancers and debilitating cardiac and dermal pathologies (27–31).

**FIGURE 1.**
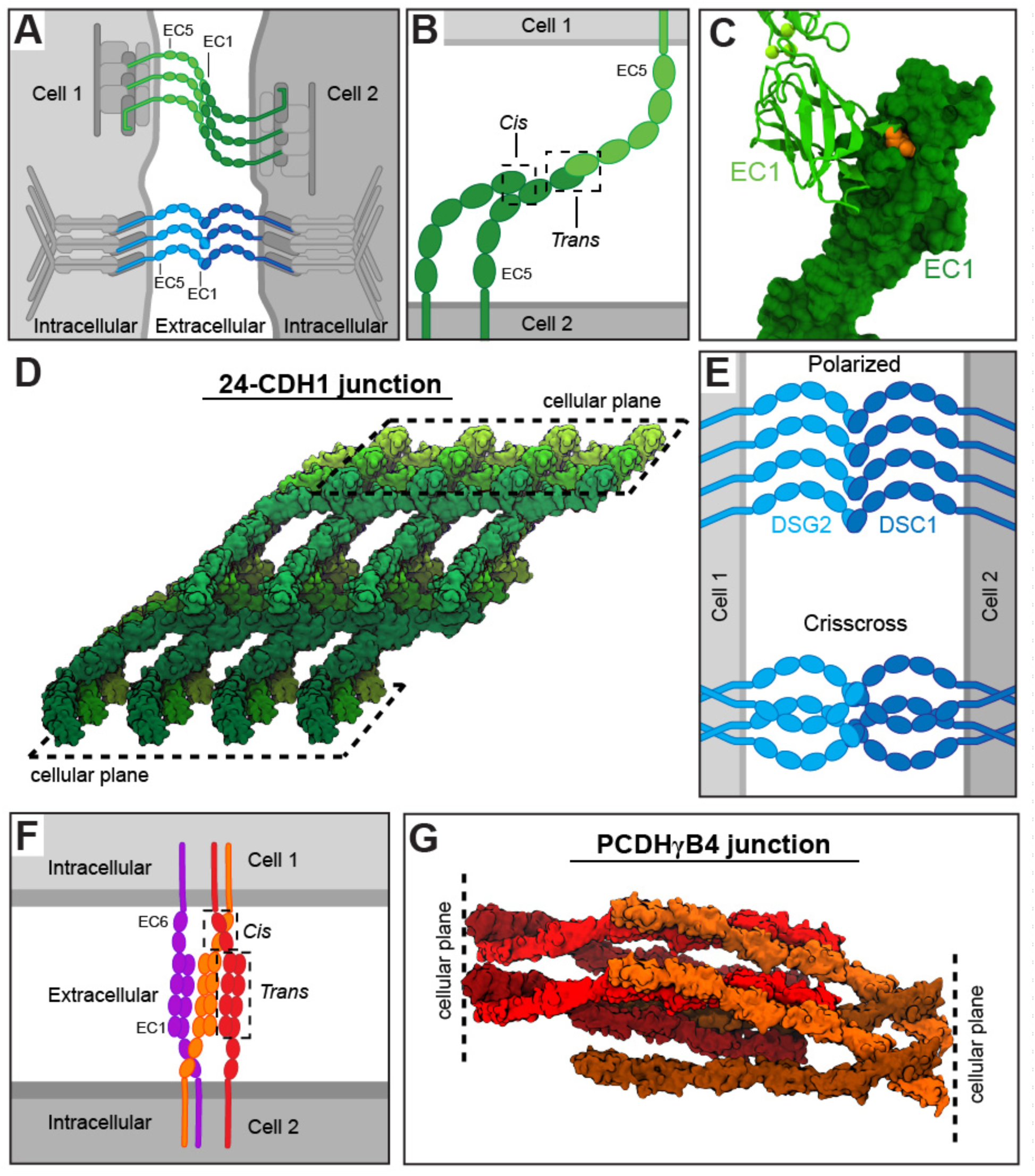
Cadherin junctions and models. (*A*) Schematic of the adherens junction (CDH1: greens) and the desmosome (DSGs and DSCs: blues) in epithelial cells. Proteins that link the cadherins to the cytoskeleton are shown in grays. (*B*) Illustration of binding modes in CDH1. *Trans*-binding occurs between EC1s from opposing cells. *Cis*-binding occurs between EC1 and EC2 of CDH1 monomers from the same cell. (*C*) Detail of the *trans* tryptophan exchange mechanism in classical cadherins. One monomer is shown in surface while the other is shown in ribbon representation. Tryptophan residue at position two (Trp^2^) in one monomer is shown in orange. (*D*) The 24-CDH1 junction shown in surface representation. The hypothetical cell surfaces are defined by the C-termini of CDH1 monomers. (*E*) Hypothetical arrangements of DSGs (light blue) and DSCs (blue) within the desmosome. The first model is a polarized arrangement in which molecules are segregated to opposite cells with their curved ectodomains aligned in the same direction. The second model is the crisscross arrangement in which DSGs and DSCs are also segregated but their curved ectodomains are rotated 180° between each dimer pair. (*F*) Schematic of clustered PCDH junction involved in neuronal self-avoidance and non-self recognition. (*G*) A model of the PCDHγB4 junction with eight monomers shown in surface representation. Hypothetical cell surfaces are defined by the C-termini of the PCDHγB4 monomers.

CDH1, DSG, and DSC proteins are all classified as classical cadherins (32–34), a subfamily that is characterized by several distinct features: the presence of an ectodomain composed of five extracellular cadherin (EC) repeats, a single helical transmembrane domain, and an intracellular domain that interacts with linker proteins and the cytoskeleton to modulate adhesion and signaling (Fig. 1, *A* and *B*) (35). Contacts between cells formed by two classical cadherins are formed through a “strand-swap” mechanism, in which a tryptophan (Trp^2^) residue at the N-terminus of one protein is inserted into a hydrophobic pocket in the N-terminal EC repeat of the other protein, and vice-versa (Fig. 1 *C*) (33, 36, 37). In cell-cell junctions composed of classical cadherins, the binding of ectodomains from opposing cells forms an array of such contacts to provide robust adhesion (38–42).

The adherens junction is believed to be formed by a well-ordered array of CDH1 monomers in a two-dimensional lattice in which both *trans* interactions between ectodomains from opposing cells and *cis* interactions between ectodomains stemming from the same cell are present (Fig. 1 *B* and *D*) (40). Evidence for this model comes from X-ray crystallography, electron microscopy data, and cell imaging (40, 43–45). Although the solution binding affinity of individual CDH1 *trans* dimers indicates a weak interaction (∼ 160 µM at 37°C) and *cis-*interactions are even weaker (∼ 1 mM), the adhesive strength of the adherens junction comes from the tight packing and collective behavior of a large number of these monomers interacting on the surfaces of adjacent cells (40, 46, 47). Experimental approaches are particularly good at describing the binding strength and affinity of individual CDH1 pairs (40, 47–53), but physical experiments that probe the mechanics of isolated cadherin lattices are difficult to perform with the proper *trans* and *cis* interactions intact (26, 54–59). The molecular mechanisms behind the mechanical response of the adherens junction have yet to be fully understood.

Compared to the adherens junction, high-resolution structural information about desmosomes is scarcer. Humans have four DSG isoforms (DSG1-4) and three DSC isoforms (DSC1-3) (41, 60). The expression levels of these differing isoforms vary spatially throughout stratified epithelia, suggesting preferential interactions depending on the specific layer of the epidermis in which the desmosome is found (5, 25). While it is known that both DSG and DSC proteins are required for the formation of the mature desmosome (61, 62), and that DSG and DSC proteins engage in homophilic and heterophilic interactions with affinities that range from ∼ 3 µM to ∼ 50 µM (41), the stoichiometric relationship between DSG and DSC proteins, whether the junction exhibits any polarity, and even the overall structural arrangements of molecules in the desmosome remain to be decisively elucidated (Fig. 1 *E*). Models from cryo-electron tomographic (cryo-ET) imaging and those based on the structure of CDH1 lattices have been suggested, and with the deposition of high-resolution crystal structures of DSG and DSC proteins these models have been further refined (38–42, 60, 63–65), yet details about their mechanical response and rupture are still unknown.

In contrast to epithelial and cardiac tissues (8, 9, 18–20, 10–17), neuronal cells are not necessarily expected to be under continuous mechanical stress. However, axon growth and neuronal circuit formation during brain development is a dynamic process in which mechanical forces play a role (66), and force-generating and load-bearing proteins are also thought to regulate synapse development and function (67–69), while action potentials can cause neuronal deformation (70–75), and traumatic brain injury can disrupt neuronal adhesion (76). Interestingly, the clustered protocadherin (PCDH) proteins, a subtype of cadherins expressed predominantly in the nervous system (77–83), form adhesive junctions (84–90) thought to signal for self-avoidance (91–93) and that may also be involved in synapse maturation and function (94–98). The clustered PCDH junctions may need to withstand mechanical stimuli, albeit in a different cellular and functional context than classical cadherins. Remarkably, the ectodomains of clustered PCDH proteins, with six EC repeats, overlap to form large adhesive antiparallel *trans* EC1-4 interfaces (Fig. 1 *F* and (*G*) (79, 84, 87, 99, 100). Analytical ultracentrifugation analyses of clustered PCDHs show that homophilic binding affinities range from ∼ 0.1 to ∼ 147 µM (85, 88, 89), and binding affinities for *cis* dimerization range from ∼ 8 to ∼ 80 µM (89). Therefore, all biophysical and structural data suggest that clustered PCDH adhesive junctions are tighter and mechanically stronger than those formed by the smaller and weaker EC1-EC1 contacts of classical cadherins, yet the mechanical properties of clustered PCDH junctions are unknown.

In a companion manuscript we explore the elasticity of single classical cadherin and clustered PCDH *trans* dimers, relevant for adhesive complex formation and for the interpretation of single-molecule force spectroscopy data. Here we focus on the mechanics of large complexes with multiple cadherin dimers involved in *trans* and *cis* interactions. We present molecular models of minimalistic cadherin-based junctions and report on simulations that explore their collective behavior and responses to tensile mechanical force. Models for the CDH1 adherens junction, two desmosome models made of DSG2 and DSC1 dimers, and a clustered PCDH junction were constructed using high resolution crystal structures (40, 41, 99) and assembled based on crystallographic lattices (CDH1; PCDHγB4) and cryo-ET maps (DSG2 and DSC1) (39, 63, 64). Steered molecular dynamics (SMD) simulations (101–104), in which C-terminal ends of monomers within complexes were stretched in opposite directions, revealed a favored shearing pathway for the adherens junction model and a resilient, two-phased tensile elastic response for the adherens junction and the desmosome systems. As observed for single dimers, curved ectodomains of classical cadherins unbend softly first, and then become stiff with unbinding proceeding without unfolding in response to tensile forces. The PCDHγB4 dimers within the junction, on the other hand, are brittle and stiff, with little extension before unbinding at force peaks that are higher than those predicted for classical cadherins. Formation of transient PCDHγB4 intermediate states adds a layer of complexity to this response. In all three junctions we observe how dimer responses within the lattices to tensile force are not equal as *cis* interactions modulate elasticity. We propose that these collective, mechanically distinct responses, modulated by *cis* contacts, are suited for function in tissue mechanics (elastic response) and self-avoidance (brittle response).

## MATERIALS AND METHODS

### Simulated Systems

Five molecular systems were built for simulation using VMD (systems S1 to S5 in Table 1) (105). All five systems had hydrogen atoms automatically added to proteins with the psfgen plugin. The protonation states of histidine residues were determined by the formation of evident hydrogen bonding partners. Residues Glu and Asp were assigned a negative charge while Lys and Arg residues were assigned a positive charge, and N-termini were assumed charged. Glycosylation sugars, alternative conformations, and crystallization reagents were not included in the models. The solvate plugin of VMD was used to add TIP3P water molecules, and the autoionize plugin was used to neutralize and randomly place ions in each system to a concentration of 150 mM NaCl.

**Table 1.** Summary of cadherin simulations.

| Label | System | $t_{\text{sim}}$<br>(ns) | Type | Start | Speed<br>(nm/ns) | Average Peak<br>Force (pN) | Size<br>(#atoms) | Initial Size (nm<sup>3</sup>) |
| --- | --- | --- | --- | --- | --- | --- | --- | --- |
| S1a | 24 CDH1 | 21.2 | EQ<sup>a</sup> | — | — | — | 3,703,940 | 66.6 x 19.7 x 29.4 |
| S1b | separation | 3.5 | SMD | S1a | 10 | 839.1 ± 115.7 |  |  |
| S1c | & relaxation | 3.5 | SMD | S1a<sup>†</sup> | 10 | 838.7 ± 125.2 |  |  |
| S1d |  | 3.5 | SMD | S1a<sup>‡</sup> | 10 | 839.1 ± 111.5 |  |  |
| S1e |  | 26.6 | SMD | S1a | 1 | 471.2 ± 64.0 |  |  |
| S1f |  | 28.0 | SMD | S1a<sup>†</sup> | 1 | 466.3 ± 120.9 |  |  |
| S1g |  | 28.5 | SMD | S1a<sup>‡</sup> | 1 | 474.0 ± 110.0 |  |  |
| S1h |  | 49.0 | SMD | S1a | 0.5 | 447.8 ± 88.3 |  |  |
| S1i |  | 50.0 | SMD | S1a<sup>†</sup> | 0.5 | 454.5 ± 64.6 |  |  |
| S1j |  | 51.2 | SMD | S1a<sup>‡</sup> | 0.5 | 458.9 ± 65.1 |  |  |
| S1k |  | 21.0 | EQ<sup>a</sup> | S1e<sup>€</sup> | — | — |  |  |
| S1l |  | 21.0 | EQ<sup>a</sup> | S1e<sup>∇</sup> | — | — |  |  |
| S2a | 16 CDH1 | 21.2 | EQ<sup>a</sup> | — | — | — | 3,102,869 | 53.0 x 20.6 x 28.0 |
| S2b | shearing | 8.0 | SMD<sup>R</sup> | S2a | 10 | 775.5 ± 97.0 |  |  |
| S2c |  | 3.5 | SMD<sup>L</sup> | S2a | 10 | 827.8 ± 138.2 |  |  |
| S2d |  | 72.7 | SMD<sup>R</sup> | S2a | 1 | 521.2 ± 114.7 |  |  |
| S2e |  | 21.0 | SMD<sup>L</sup> | S2a | 1 | 499.9 ± 98.4 |  |  |
| S2f |  | 141.6 | SMD<sup>R</sup> | S2a | 0.5 | 474.7 ± 82.4 |  |  |
| S2g |  | 38.3 | SMD<sup>L</sup> | S2a | 0.5 | 427.6 ± 87.8 |  |  |
| S3a | DSG2-DSC1 | 21.2 | EQ<sup>b</sup> | — | — | — | 1,882,422 | 54.5 x 45.2 x 50.6 |
| S3b | polarized | 4.0 | SMD | S3a | 10 | 733.7 ± 163.3 |  |  |
| S3c |  | 30.3 | SMD | S3a | 1 | 373.4 ± 75.9 |  |  |
| S3d |  | 270.4 | SMD | S3a | 0.1 | 347.1 ± 64.1 |  |  |
| S4a | DSG2-DSC1 | 22.4 | EQ<sup>b</sup> | — | — | — | 3,485,695 | 63.7 x 26.0 x 20.7 |
| S4b | crisscross | 3.7 | SMD | S4a | 10 | 884.3 ± 128.2 |  |  |
| S4c |  | 19.8 | SMD | S4a | 1 | 499.7 ± 83.9 |  |  |
| S5a | PCDHγB4 | 21.2 | EQ<sup>a</sup> | — | — | — | 3,133,122 | 74.9 x 26.6 x 16.2 |
| S5b |  | 4.3 | SMD | S5a | 10 | 1293.7 ± 301.5 |  |  |
| S5c |  | 14.1 | SMD | S5a | 1 | 943.7 ± 309.0 |  |  |
| S5d |  | 103.4 | SMD | S5a | 0.1 | 607.9 ± 146.3 |  |  |
| **Total** |  | 1128.1 |  |  |  |  |  |  |

<sup>†</sup> denotes that simulations started from simulation S1a at 16.2 ns.
<sup>‡</sup> denotes that simulations started from simulation S1a at 19.2 ns.
<sup>€</sup> denotes that simulation S1k started from simulation S1e at 21.1 ns.
<sup>∇</sup> denotes that simulation S1l started from simulation S1e at 23.1 ns.
<sup>a</sup> denotes constraints were used on the C-terminal Ca atoms during equilibration
<sup>b</sup> denotes constraints were not used on the C-terminal Ca atoms during equilibration
<sup>R</sup> denotes the 16-CDH1 junction was sheared against the natural tilt of CDH1
<sup>L</sup> denotes the 16-CDH1 junction was sheared along the natural tilt of CDH1

The first two molecular systems represented minimalistic adherens junctions and were constructed using either 24 or 16 monomers from the crystal lattice of the mouse CDH1 homodimer structure (PDB code: 3Q2V) (40). Coordinates for symmetry monomers in the lattice were obtained using COOT (106). Missing residues were added by substituting known residue coordinates from another chain in the asymmetric unit of the crystal where those residues were present. The larger CDH1 junction with 24 monomers was built so that the 12 dimers at different locations within the lattice would have duplicates that shared similar *trans*- and *cis*-dimerization profiles within the same system. To reduce overall system size and to optimize use of computational resources, we simulated this system in a rotated fashion (Fig. S1 *A* and *B* in the Supporting Material). The proteins were first aligned along a primary axis and solvated, and then the solvated system was rotated by a series of right-handed basic rotations around their axes (*θ_y_* = 45° and *θ_z_* = 50°) to align the hypothetical cellular planes perpendicular to the *x*-axis (Fig. S1 *A*, left panel). This setup allowed stretched CDH1 monomers along the *x*-axis to move into space vacated in the neighboring periodic cell (Fig. S1 *B*). The smaller CDH1 junction with 16 monomers (8 dimers) was built to study shearing and was not rotated.

The third and fourth systems represented minimalistic desmosomes and were built using the structures of human DSG2 EC1-5 (PDB code: 5ERD) and human DSC1 EC1-5 (PDB code: 5IRY) (41). These were selected for simulation because of their high resolution and their completeness as both structures showed the entire ectodomains of DSG2 and DSC1. The heterodimer of DSG2-DSC1 (Fig. S2 *A*) was created as described in the companion paper and used to assemble a desmosomal junction with 8 monomers (4 dimers). To build the first model of the desmosome junction we used the crystallographic lattice of C-cadherin (PDB code: 1L3W) (37) as a template. The first 400 atoms of either DSG2 or DSC1 were aligned to the first 400 atoms of C-cadherin in a conformation in which all DSG2 monomers were on one side of the lattice and all DSC1 on the other in a polarized fashion (Fig. S2 *B*). Clashing was relaxed through minimization with backbone constraints in vacuum and further relaxation was achieved through minimization and equilibration after solvation and ionization. This “polarized” DSG2-DSC1 system was rotated by *θ_y_* = 140° and *θ_z_* = 35° after solvation to align the hypothetical cellular planes perpendicular to the *x*-axis and to allow for stretching along this axis as done for the large CDH1 adherens junction system.

The fourth system (second desmosomal system) was created based on the model proposed by Al-Amoudi *et al*. (39) in which the C-cadherin structure was fit to a three-dimensional cryo-ET map of an intact human epidermis desmosome, but using our DSG2-DSC1 heterodimer model based on the most recent structural data for these proteins (41). Thus, we first used VMD to align the DSG2-DSC1 heterodimer to one of the EC1 monomers in the structure of N-cadherin (PDB code: 1NCH) (36), which has two antiparallel monomers in the asymmetric unit and was used as a reference in (39). A second DSG2-DSC1 heterodimer was aligned to the other EC1 monomer in the structure. The resulting model, containing two DSG2-DSC1 dimers, was manually and slightly translated to better match the geometry seen in the reconstructed desmosomal cryo-ET density (39, 64). Two of these models were aligned to create a system with 8 monomers. The resulting system features a “crisscross” structure, in which the orientation of each adjacent dimer is rotated by 180° around an axis normal to the cell plane (Fig. S2 *C*), as opposed to the lattice built based on C-cadherin, in which every dimer is assembled in the same orientation (Fig. S2 *B*). This crisscross DSG2-DSC1 system was not rotated. Attempts to build another junction conformation in which the orientation of DSG2-DSC1 heterodimers was alternated in a “checkerboard” pattern were unsuccessful as we were not able to resolve clashes among monomers (Fig. S2 *D*).

The last, fifth system, representing a clustered PCDH junction, was built using the crystal structure of mouse PCDHγB4 reported by Brasch *et al*. (PDB code: 6E6B) (99). Residues that were missing from one of the monomers in the structure (residue 253 to 258) were modeled using the CHARMM-GUI (107) and added to the original structure. The system contains 8 monomers of PCDHγB4 extracted from the crystallographic lattice with COOT and features three *trans* interactions and four *cis* interactions. Two PCDHγB4 monomers at the edges of the junction lack a *trans*-binding partner but do have *cis* interactions. The PCDHγB4 system was rotated by *θ_y_* = 22° and *θ_z_* = 10° after solvation to allow for stretching along the *x*-axis.

### Simulations

Molecular dynamics (MD) simulations using explicit solvent (108–116) were carried out using NAMD 2.11, 2.12, and 2.13 (117) utilizing the CHARMM36 (118) force field for proteins with the CMAP (119) backbone correction. Simulations of the 24-CDH1 junction used GPU acceleration (120). A switching distance of 10 Å with a cutoff of 12 Å was used for van der Waals interactions with pair list generation within 13.5 Å updated every 40 fs. To compute long-range electrostatic forces, the particle mesh Ewald (PME) method (121) with a grid point density of > 0.5 Å^-3^ was used for the CDH1 and PCDH systems and a grid point density of > 1 Å^-3^ was used for the other systems. A uniform integration time step of 2 fs for evaluation of bonded and non-bonded interactions was used together with the SHAKE algorithm (122). Constant temperature (*T* = 300 K) was maintained using Langevin dynamics with a damping coefficient of γ = 0.1 ps^-1^ unless otherwise stated. The hybrid Nosé-Hoover Langevin piston method with a 200 fs decay period and a 50 fs damping time constant was used to maintain constant number, pressure, and temperature simulation conditions (*NpT*) at 1 atmosphere (117). Constraints on C_α_ atoms were applied using a harmonic potential with a spring constant of *k_c_ =* 1 kcal mol^-1^ Å^-2^. All systems were minimized for 5000 steps followed by a backbone-constrained equilibration for 200 ps, a 1 ns bridging equilibration with a Langevin damping coefficient of γ = 1 ps^-1^, and a final equilibration of 20 ns with γ = 0.1 ps^-1^ and constraints applied only on the C-terminal C_α_ atom in each monomer for the CDH1 and PCDH junctions and no constraints for the desmosomal junctions.

SMD simulations were carried out using the NAMD Tcl forces interface to implement constant-velocity stretching (101–104). Independent virtual springs (stiffness *k_s_* = 1 kcal mol^-1^ Å^-2^) were attached to each C-terminal C_α_ atom, and the free ends of these springs were moved at a constant velocity along the *x*-axis and away from the protein (tensile mode). Unlike slab schemes (123–125), forces are not re-distributed as bonds rupture. Additional harmonic constraints in the *y* and *z* directions were applied to guide stretching along the *x*-axis only. For each system, SMD simulations were carried out at constant velocities of 10, 1, and either 0.5 or 0.1 nm/ns. Shearing simulations were carried out similarly by moving the free ends of the springs in the indicated directions and adding harmonic constraints in the other perpendicular directions. The applied forces, calculated from the extension of the virtual springs and from harmonic constraints, were recorded every 40 fs. Whole system coordinates were saved every 1 ps.

### Simulation analysis procedures and tools

Force plots include the magnitude of the total applied force from virtual springs and harmonic constraints on the C-terminal C_α_ atoms of each *trans* dimer pair. We computed end-to-end distances for complexes as the magnitude of the distance between C-terminal C_α_ atoms, unless otherwise stated. Stiffness for complexes was computed using linear regression fits of the force versus distance plots. A 50-ps running average, to eliminate local fluctuations, was used to obtain maximum force peaks. Buried surface area (BSA) was computed in VMD by measuring solvent accessible surface area (SASA) for individual monomers and by subtracting SASA for the complex, with BSA for interacting molecules A and B computed as BSA_AB_ = ½ (SASA_A_ + SASA_B_ - SASA_AB_). We used Xmgrace to generate plots and the molecular graphics program VMD (105) to analyze trajectories, render molecular images, and create videos.

## RESULTS

Simulations of individual cadherin dimeric complexes presented in a companion manuscript revealed a significant difference in the mechanical responses of classical cadherins versus clustered PCDH proteins. Applied forces to CDH1 and desmosomal complexes (DSG2-DSG2, DSG2-DSC1, DSC1-DSC1) resulted in soft unbending over ∼ 10 nm, followed by stiffening and unbinding without unfolding. On the other hand, clustered PCDH dimeric complexes (PCDHα7, PCDHβ6, PCDHγB3) lacked the soft unbending phase and unbound at larger forces, suggesting a brittle response that contrasts with the mechanically resilient behavior of classical cadherins. To understand the behavior of classical cadherins and clustered PCDH proteins in the context of larger complexes where lateral *cis* contacts are relevant, and to determine whether the collective behavior is different than the response of individual dimers, we built and simulated five systems representing different types of adhesive junctions. We describe below results from equilibrium and SMD simulations for each of these systems, including for two models representing adherens junctions, two models representing desmosomes, and one representing a clustered PCDH junction.

### Elastic Mechanical Response of Adherens Junction Models

Adherens junctions are expected to be under tension generated by the actomyosin cytoskeleton and may experience more dramatic tensile and shearing force challenges during tissue morphogenesis and function as well as in wounding (58, 126–131). Two similar models of adherens junction were simulated, including a first model with 24 CDH1 ectodomains (∼ 3.7-M atom system) used to study the response of the junction to tensile stretching forces and a second model with 16 CDH1 ectodomains (∼ 3.1-M atom system; see Materials and Methods and Table 1) used to study the response of the junction to shearing forces. Each model was equilibrated before forces were applied to either stretch or shear the systems as described below.

#### Soft unbending and unbinding with location-dependent mechanical responses during tensile stretching

The first model representing a minimalistic adherens junction has the entire ectodomains of 24 CDH1 monomers forming 12 *trans* dimers at positions that we labeled P01-P02, P03-P04, …, P23-P24, with even numbers for monomers at one side of the junction and odd numbers for monomers at the other side (Figs. 1 *A* and 2, *A* and *D*). This system was equilibrated for 20 ns (simulation S1a) with harmonic constraints applied to the C-terminal C_α_ atoms to mimic attachment to the underlying cytoskeleton. During the equilibration we observed stable *trans* dimers as well as curvature changes across the monomers, with some compressing and others expanding to compensate and thus satisfy the imposed constraints without rupturing of interfaces. Conformations obtained throughout the equilibration were used as starting points for SMD simulations at stretching speeds of 10 nm/ns, 1 nm/ns, and 0.5 nm/ns, each done in triplicates (simulations S1b-j; Videos S1 and S2). In each of the SMD simulations, C-termini from monomers in opposite sides of the junction were moved in opposite directions (tensile mode) to induce stretching and unbinding, with harmonic constraints applied to the C-terminal C_α_ atoms in the plane perpendicular to the stretching axis to mimic attachment to cytoskeletal elements and with forces applied to each monomer recorded to monitor their mechanical response (Fig. 2 *A-C*). Below we discuss results for the slowest stretching speed simulations as good representatives of all the SMD simulations for this system.

**FIGURE 2.**
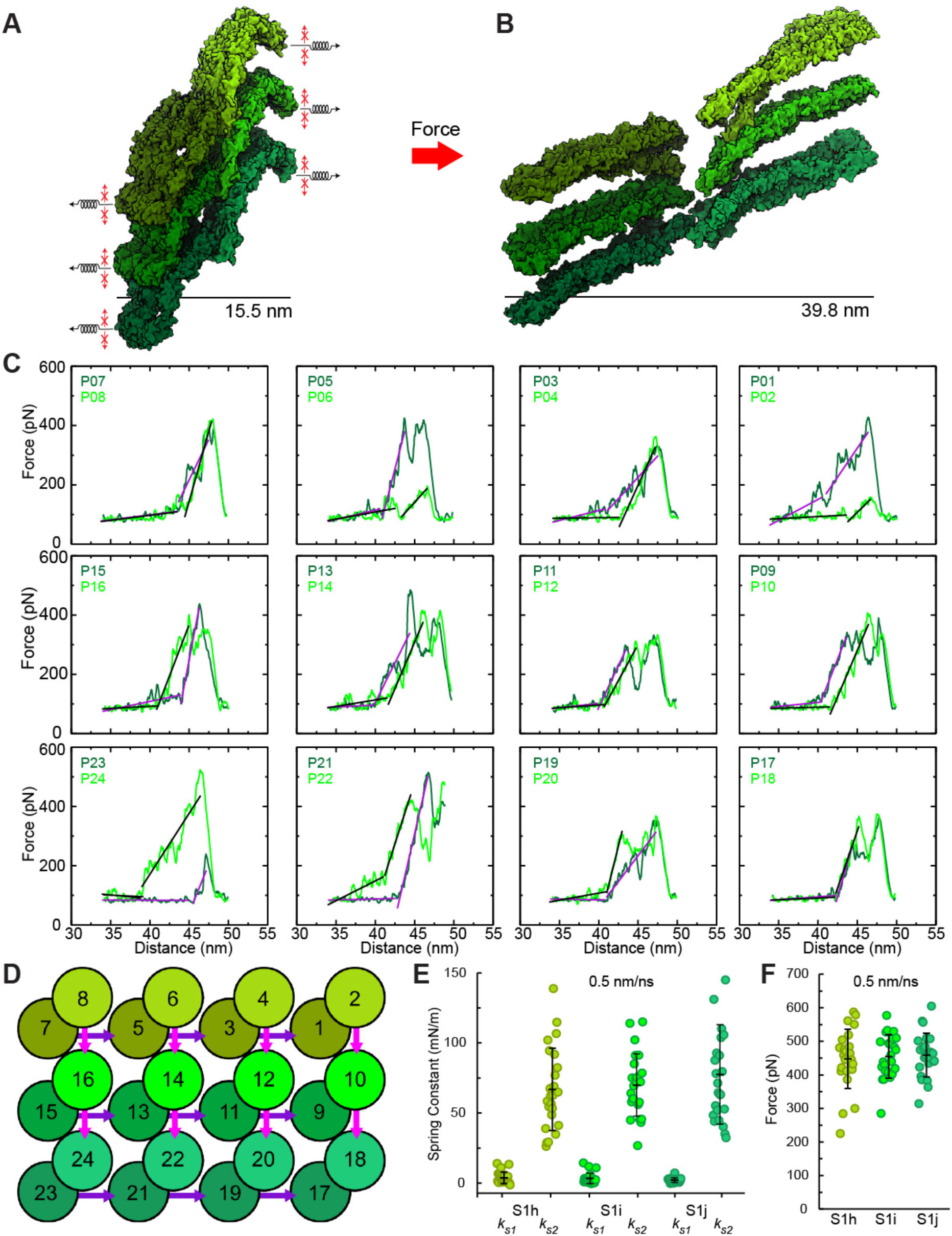
Forced unbinding of 24-CDH1 adherens junction. (*A*) Snapshot of the initial state of the CDH1 junction. Springs with arrows show the SMD pulling direction. Small red arrows indicate constraints in the plane applied to mimic cytoskeleton attachment. Black bar is the distance between hypothetical cellular planes. (*B*) The stretched and ruptured state of the 24-CDH1 junction (simulation S1h at 0.5 nm/ns; Table 1). (*C*) Force versus end-to-end distance plots for constant-velocity stretching of individual CDH1 *trans* dimer pairs within the junction (0.5 nm/ns, simulation S1h). Bright green lines are for CDH1 monomers on the right side of the junction in (*A*) and (*B*) or the top side of the junction in (*D*); dark green lines are for CDH1 monomers on the left side in (*A*) and (*B*) or the bottom side in (*D*); purple and black lines are linear fits used to determine elasticity. Plots are arranged to reflect the position of CDH1 *trans* dimers within the junction, as labeled in (*D*) (P01 – P24). (*D*) Schematic of the 24-CDH1 junction. Positions of CDH1 monomers are labeled P01 – P24. *Trans* interactions are represented by an overlap of circles. Arrows indicate *cis* interactions: Pink arrows are *cis* interactions among top layer monomers while purple arrows are for *cis* interactions among bottom layer monomers. Arrows show *cis*-interaction directionality with their base representing EC1 of the originating CDH1 monomer and their arrowhead representing the binding surface of EC2 on another CDH1 monomer. (*E*) Spring constants for phase 1 (*k_s_*_1_) and phase 2 (*k_s_*_2_) from simulations S1h-j. Values for individual monomers indicated as circles. Average and standard deviation shown as black bars (olive circles - simulation S1h; bright green circles - simulation S1i; forest green circles - simulation S1j). (*F*) Peak force required to rupture *trans* dimers in the 24-CDH1 junction. Circles and bars as in (*E*).

The 24-CDH1 junction stretched at 0.5 nm/ns displayed an initial elongation phase in which CDH1 *trans* dimers unbent at small forces (∼ 50 to 150 pN) over extensions of ∼ 5 to 10 nm and with a soft effective spring constants of *k_s1_* = 3.7 ± 4.1 mN/m for simulation S1h, *k_s1_* = 3.3 ± 3.8 mN/m for simulation S1i, and *k_s1_* = 2.1 ± 1.6 mN/m for simulation S1j (Figs. 2 *E* and S3 *A*; averages computed from slopes of force *versus* end-to-end distance curves on each side of the junction for all *trans* dimers). Computing these spring constants using the separation of the membrane planes instead yields *k_s1_* = 1.9 ± 2.2 mN/m for simulation S1h, *k_s1_* = 1.8 ± 2.3 mN/m for simulation S1i, and *k_s1_* = 1.1 ± 1.0 mN/m for simulation S1j (Fig. S3 *A*). In some cases we observed flat or even negative slopes in the force versus end-to-end distance curves for some dimers. We attribute this to compressed states that emerged during equilibration, and to *cis* interactions that communicated force from a neighboring monomer thus inducing extension without the need of applied force to that particular CDH1 monomer.

After this initial soft extension mediated by unbending, the CDH1 *trans* dimers became stiffer, with effective spring constants of *k_s2_* = 66.9 ± 29.4 mN/m for simulation S1h, *k_s2_* = 69.9 ± 22.3 mN/m for simulation S1i, and *k_s2_* = 77.5 ± 35.6 mN/m for simulation S1j (Fig. 2 *E*). Softer values are obtained when using the separation of membrane planes with effective spring constants of *k_s2_* = 46.3 ± 20.0 mN/m for simulation S1h, *k_s2_* = 47.9 ± 14.6 mN/m for simulation S1i, and *k_s2_* = 53.1 ± 24.7 mN/m for simulation S1j (Fig. S3 *B*). Compared to simulations of isolated CDH1 *trans* dimers, the *trans* dimers within the junction have a similar initial soft elastic response to force, but spring constants associated with the stiffer phase before unbinding are larger, which indicates that CDH1 *cis* interactions may increase the adherens junction stiffness.

After unbending and stretching, unbinding for each of the *trans* dimers proceeded without unfolding when swapped Trp^2^ residues dislodged and other contacts broke at the maximum force peak of *F*_p_ ∼ 447.8 ± 88.3 pN for simulation S1h, *F*_p_ ∼ 454.5 ± 64.6 pN for simulation S1i, and *F*_p_ ∼ 458.9 ± 65.1 pN for simulation S1j (Fig. 2 F; averages computed over maximum force peaks monitored across all *trans* dimers on each side of the junction). Disulfide bonds at the EC5 C-termini prevented unraveling, and rupture of individual *cis* and *trans* interactions manifested in various force peaks for the CDH1 *trans* dimers within the junction (Fig. 2 *C*, Fig. S4). The average force value of the first discernable peak is *F*_p1_ ∼ 375.2 ± 112.7 pN at an end-to-end distance of ∼ 44.1 ± 1.7 nm (membrane-to-membrane distance of ∼ 32.4 ± 2.5 nm) for simulation S1h, slightly smaller than the maximum force peak at an end-to-end distance of ∼ 46.8 ± 0.9 nm (membrane-to-membrane distance of ∼ 36.1 ± 1.3 nm; Fig. S3 *C*). We monitored BSA for *trans* and *cis* dimers and found a good correlation between *trans* BSA decrease and the final force peak for individual monomers at the slowest stretching speed (simulation S1h; Fig. S5). The loss of *cis* interactions reported by a drop in *cis* BSA did not always correlate well with force peaks throughout the junction (Fig. S5). The *cis* contacts for a monomer typically separated first, before unbinding, regardless of the number of *cis* contacts for the monomer at a given position within the junction. However, *cis* contacts did not always rupture and, in a few cases, re-formed after *trans* unbinding (Fig. S5).

The mechanical responses of CDH1 *trans* dimers, monitored through force *versus* end-to-end distance curves, were dependent on their location within the junction. For instance, we monitored similar force profiles for dimers at two corners of the 24 CDH1 junction, corresponding to positions P01-P02 and P23-P24. However, the force profiles had swapped curves, as the force curve for the monomer in position P01 was similar to the force curve for the monomer in position P24 at the other side of the junction (simulations S1h-S1j; Figs. 2 *C* and *D* and S4). This swapping of curves was also observed for monomers at positions P02 and P23. The swapped curves reflect the equivalent type of *cis* contacts shared by the CDH1 *trans* dimers at positions P01-P02 and P23-P24 (Fig. 2 *D*). In contrast, CDH1 monomers in the opposite corners at positions P07-P08 and P17-P18 do not share equivalent *cis* contacts and therefore their force profiles are different from each other and from the force profiles observed for monomers at P01-P02 and P23-P24 (simulations S1h-S1j; Fig. 2, *C* and *D*). The force profiles for CDH1 *trans* dimers at the edges of the junction and center of the junction were more complex because of their increased number of *cis* interactions (simulations S1h-S1j; Figs. 2, *C* and *D* and S4). Overall, despite location-dependent force profiles for various *trans* dimers that may reflect the collective influence of *cis* contacts within the junction, we always observed a two-phased mechanical response with soft unbending and unbinding without unfolding at the slowest stretching speed used for this first CDH1 system.

#### Fast recovery of junction architecture after limited tensile stretching

Electron microscopy images (132, 133) and simulations (134) of classical cadherin ectodomains suggest that their bent shape is an intrinsic property determined by their EC linker regions. Given that in a companion manuscript we report rapid re-bending after unbinding in simulations of individual classical cadherin *trans* dimers, we hypothesized that the stretched CDH1 junction would quickly recover its shape and architecture if C-termini of individual monomers were released and further equilibrated. We tested this hypothesis by releasing applied forces after stretching the junction while keeping constraints in the membrane plane to mimic stable connections to the underlying cytoskeleton (simulation S1e followed by S1k and S1l). Release and further relaxation simulations started either after all monomers had been straightened out but no *trans* interaction had been lost (pre-ruptured state obtained after 21.1 ns of stretching at 1 nm/ns; Fig. 3 *A*), or after three *trans* bonds (out of 12) had been ruptured and a fourth one had Trp^2^ residues dislodged from their hydrophobic pockets (partially ruptured state obtained after 23.1 ns of stretching at 1 nm/ns; Fig. 3, *D*).

**FIGURE 3.**
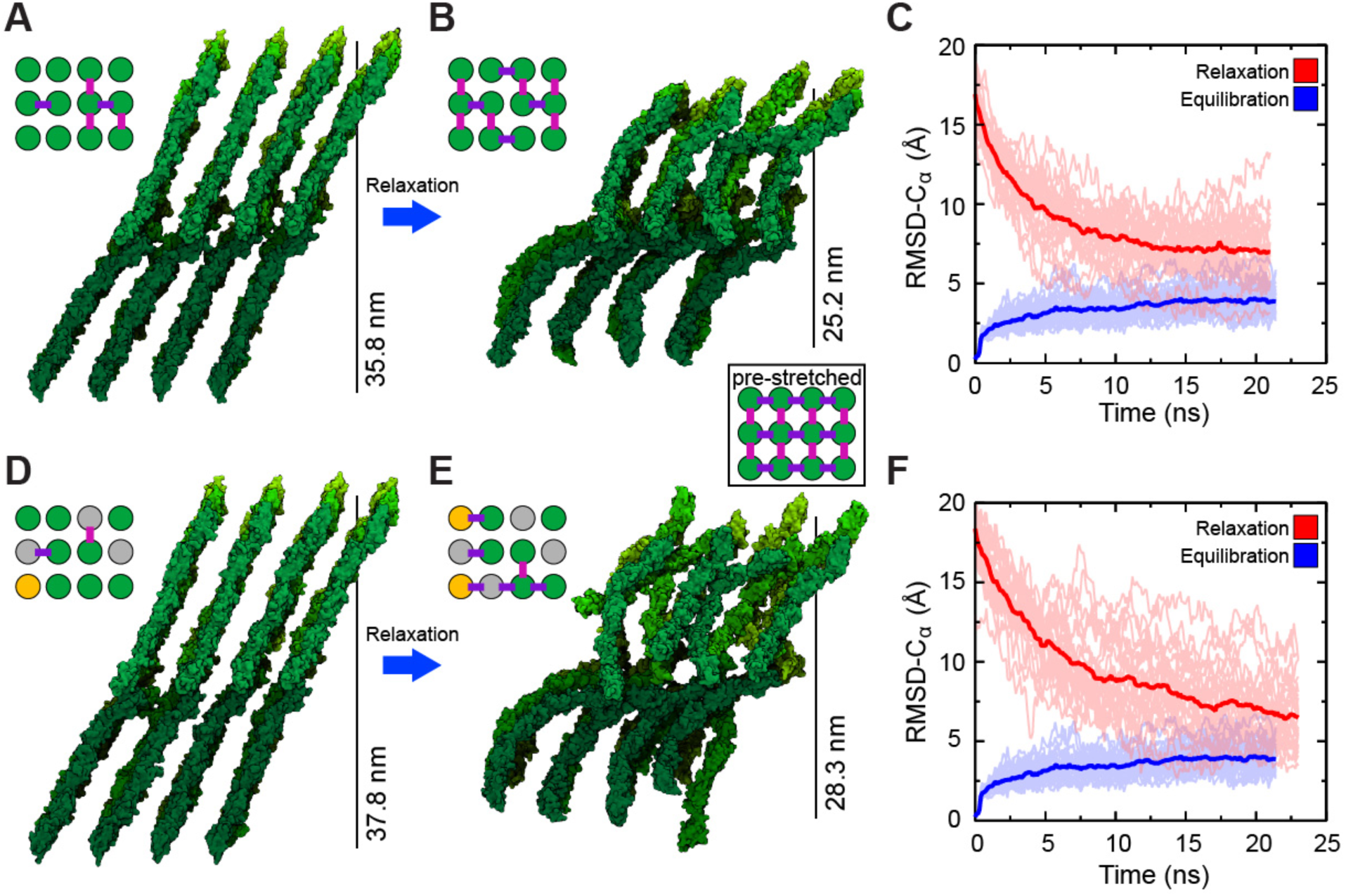
Relaxation of stretched 24-CDH1 adherens junction. (*A*) The 24-CDH1 junction after 21.1 ns of stretching at 1 nm/ns (simulation S1e; Table S1). All *trans* interactions are present as indicated by green circles in the inset. Purple and pink bars represent intact *cis* interactions on the bottom and top layer, respectively. The hypothetical distance between cellular planes is indicated. (*B*) Stretched 24-CDH1 junction in (*A*) after 21 ns of relaxation dynamics with C-terminal C_α_ atoms constrained in the plane applied to mimic cytoskeletal attachment. Individual CDH1 monomers re-bend and some *cis* contacts are re-established. Boxed inset is a schematic of the *trans* and *cis* bond prior to stretching. (*C*) RMSD-C_α_ versus time plot for the equilibration (individual monomers, pale blue; average RMSD-C_α_, blue) and for the relaxation (individual monomers, pale red; average RMSD-C_α_, red) of the 24-CDH1 junction after 21.1 ns of stretching. (*D*) The 24-CDH1 junction after 23.1 ns of stretching at 1 nm/ns (simulation S1e). Three *trans* interactions are broken as indicated by grey circles. The yellow circle indicates a *trans* interaction where the Trp^2^ residues have left their hydrophobic pockets but the CDH1 monomers have not lost contact. *Cis* interactions colored as in (*A*). (*E*) Stretched 24-CDH1 junction in *C* after 21 ns of relaxation dynamics with cytoskeletal constraints. Individual CDH1 monomers re-bend, but the original junction architecture is not recovered. Some *cis* interactions are re-formed during the simulation while some *trans* contacts are lost. (*F*) RMSD-C_α_ versus time plot for the equilibration and relaxation of the 24-CDH1 junction after 23.1 ns of stretching. Colored as in (*C*).

As expected, individual ectodomains quickly started to re-bend after relaxation was started for the pre-ruptured state of the CDH1 junction with their C-termini free to move in the stretching direction only. The CDH1 junction superficially resembled the un-stretched starting conformation of the system after 21 ns of relaxation (simulation S1k; Fig. 3 *B*; Video S3), which was also reflected in a decrease of RMSD-C_α_ for CDH1 monomers as their inherent curvature returned (Fig. 3 *C*). In contrast, 23 ns of relaxation for the partially ruptured state of the CDH1 junction resulted in a system that did not resemble the original junction, even though individual monomers recovered their curvature (simulation S1l; Fig. 3 *E* and *F*; Video S4). Re-formation of some *cis* interactions was observed in both relaxations (Fig. 3, *B* and *E*), but recovery of ruptured Trp^2^-mediated *trans* interactions did not occur and was not expected in such short timescale. These simulations indicate that the CDH1 adherens junction can quickly recover from non-rupturing stretching (∼ 10 nm) demonstrating a resilient response akin to a molecular shock absorber.

#### Preferential separation pathway during shearing

Cell-cell junctions are also expected to experience shear stress during morphogenesis or in tissues that are exposed to fluid flow (128–131). A second model representing a minimalistic adherens junction was built to study the effect of shearing forces on it. This system has the entire ectodomains of 16 CDH1 monomers forming 8 *trans* dimers at positions labeled P01-P02, P03-P04, …, P15-P16; it was equilibrated for 20 ns (simulation S2a) with harmonic constraints applied to the C-terminal C_α_ atoms to mimic attachment to the underlying cytoskeleton. As with the 24-CDH1 junction, *trans* dimers in the 16-CDH1 junction remained stable during equilibration. The final conformations obtained after equilibration were used as starting points for SMD simulations at stretching speeds of 10 nm/ns, 1 nm/ns, and 0.5 nm/ns, each done by stretching in two different directions that induced distinct separation pathways (simulations S2b-g). In each of the SMD simulations, C-termini from monomers in opposite sides of the junction were moved in opposite directions, but in one set of simulations the shearing pathway was set to stretch the CDH1 *trans* dimers along their natural tilt (Fig. 4 *A*) while in the other set the shearing pathway was set to push against it (Fig. 4 *D*). In all shearing SMD simulations we applied harmonic constraints to the C-terminal C_α_ atoms in the plane perpendicular to the stretching axis to mimic attachment to cytoskeletal elements and recorded forces applied to each monomer to monitor their mechanical response. Below we discuss results for the slowest stretching speed simulations as good representatives of all the SMD simulations for this system.

**FIGURE 4.**
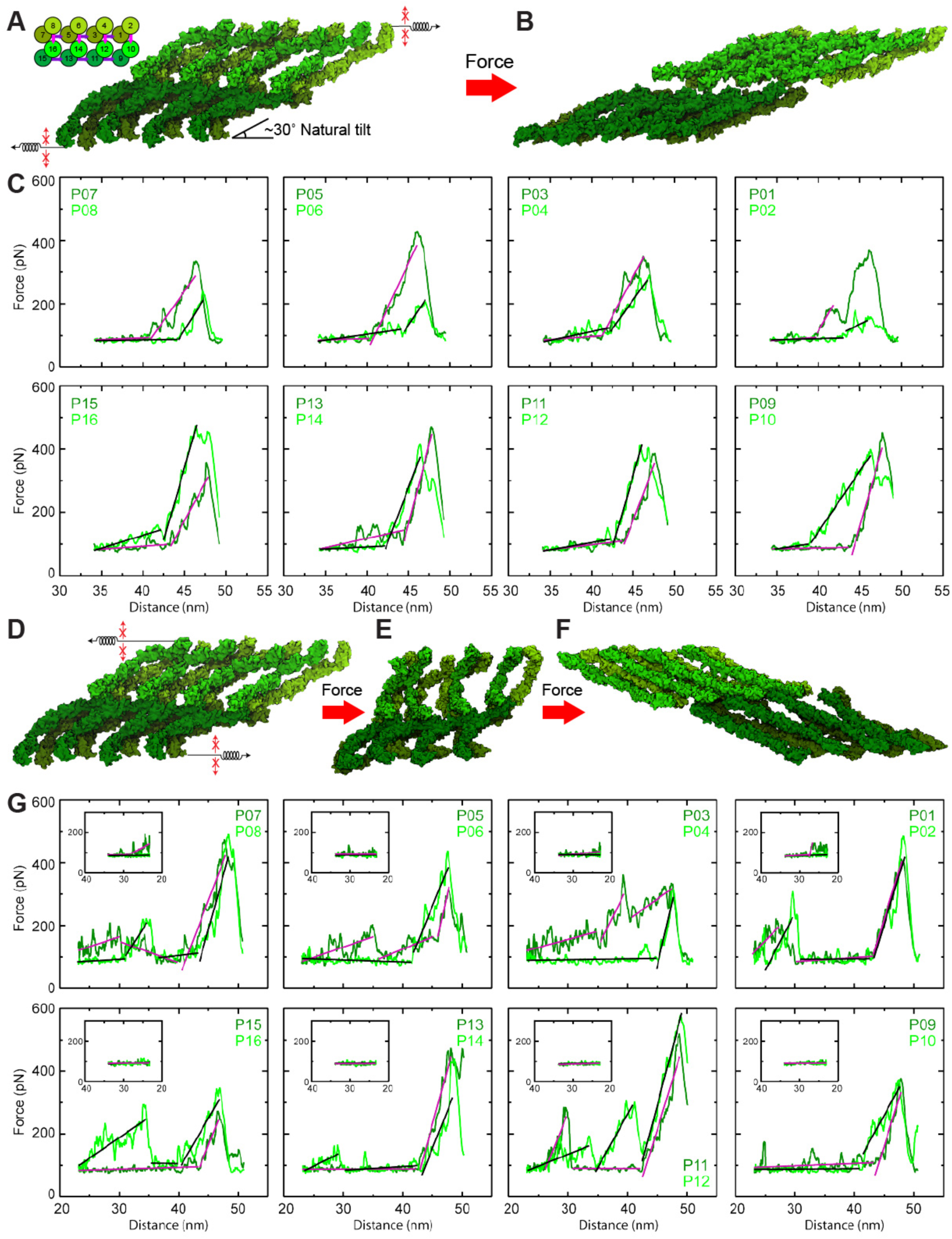
Shearing of the 16-CDH1 adherens junction. (*A*) Starting state of the 16-CDH1 junction. Springs with arrows indicate the stretching direction. Small red arrows indicate constraints in the plane applied to mimic cytoskeleton attachment. Inset schematics show the system with positions labeled P01 – P16. *Trans* interactions are represented by an overlap of circles. Arrows indicate *cis* interactions as in Fig. 2 *D*. The natural tilt of CDH1 monomers with respect to the hypothetical cellular membrane plane is indicated. (*B*) The 16-CDH1 junction after shearing at 0.5 nm/ns along the direction indicated in (*A*). There was no loss of *cis* interactions throughout the trajectory. (*C*) Force versus end-to-end distance plots for constant-velocity shearing (0.5 nm/ns) of individual CDH1 *trans* dimer pairs within the junction shown as in Fig. 2 *C* (simulation S1h). Plots are arranged to reflect the position of CDH1 *trans* dimers within the junction, as labeled in the inset in (*A*) (P01 – P16). (*D*) Starting state of the 16-CDH1 junction shown in (*A*), but with springs indicating the shearing direction opposite of the natural tilt. (*E* and *F*) Compressed and ruptured states of the 16-CDH1 junction, respectively. (*G*) Force versus end-to-end distance plots during stretching for constant velocity shearing (0.5 nm/ns) of individual CDH1 *trans* dimer pairs within the junction shown as in (*C*). Inset is the force versus end-to-end distance during compression.

When the 16-CDH1 junction was sheared along the natural tilt of CDH1 monomers, the separation of layers was fast and the force response (Fig. 4 *A – C*; Video S5) mainly consisted of two phases as described for the 24-CDH1 junction responding to tensile forces, with soft unbending of *trans* dimers followed by a stiffening of their force response before unbinding. Separation through unbinding occurred after ∼ 10 nm of end-to-end extension, and a relative displacement of ∼ 15.7 nm for the cell planes. Interestingly, when the 16-CDH1 junction was sheared in the opposite direction, we observed an initial compression of the CDH1 monomers with little resistance accompanied by the loss of *cis* interactions during or immediately after the compression of the system. A multi force peak response that reflected a longer and more cumbersome separation pathway involved a relative displacement of cell planes of ∼ 85.0 nm until unbinding of all monomers (Fig. 4 *D – G*; Video S6). The maximum force peaks upon unbinding in shearing simulations at 0.5 nm/ns were similar for both separation pathways, with values of *F*_p_ = 427.6 ± 87.8 pN and *F*_p_ = 474.7 ± 82.4 pN for simulations 2f and 2g, respectively (averages computed over forces monitored across all *trans* dimers on each side of the junction). However, the presence of multiple force peaks throughout a significantly longer separation pathway when shearing the systems against the natural tilt of CDH1 monomers indicates that individual adherens junction patches may have a preferential shearing pathway. These results suggest that the orientation of the CDH1 lattice could have functional implications for directionality in cell migration and tissue morphogenesis.

### Resilient Mechanical Response of Desmosomal Junction Models

Desmosomes in cardiac and epithelial tissues, including the skin, are expected to experience and withstand mechanical forces. However, details of how the various DSG and DSC desmosomal cadherins arrange and distribute to form mechanically strong junctions remain to be established. Two different models of minimalistic desmosome junctions were simulated to study their response to tensile stretching forces, including a polarized junction built based on the C-cadherin crystallographic lattice (1.8 M-atom system) and a crisscross junction based on the Cryo-ET data from Al-Amoudi *et al*. (39, 64) (3.4 M-atom system; see Materials and Methods and Table 1). Each model was equilibrated freely for 20 ns before force was applied to explore their mechanical response as described below.

#### Soft unbending and unbinding in a polarized desmosome during tensile stretching

The first model representing a minimalistic desmosome junction has the entire ectodomains of four DSG2 monomers forming *trans* dimers with four DSC1 monomers at positions labeled P01-P02, P03-P04, P05-P06, and P07-P08 (even numbers for DSG2 and odd numbers for DSC1 monomers) (Fig. 5 *A* and *D*). This polarized system, built based on the C-cadherin crystallographic lattice, was equilibrated for 20 ns (simulation S3a). During equilibration, the *cis* contacts present in the initial model remained stable. However, the Trp^2^ residue of DSC1 at position P03 slipped out of the hydrophobic pocket of its DSG2 *trans* binding partner at P04. Nevertheless, this *trans* interaction remained stable due to a series of specific interactions that included the DSG2-DSC1 pairs Glu^31^ - Arg^1^, Glu^30^ - Lys^97^, Lys^23^ - Glu^99^, Glu^91^ - Arg^1^, and Lys^92^ - Arg^1^ (backbone interaction). After equilibration, SMD simulations in which C-termini from monomers in opposite sides of the junction were moved in opposite directions so as to induce tensile stretching and unbinding, and were carried out at stretching speeds of 10 nm/ns, 1 nm/ns, and 0.1 nm/ns (simulations S3b-d). Additional harmonic constraints were applied to the C-terminal C_α_ atoms in the plane perpendicular to the stretching axis to mimic attachment to cytoskeletal elements. We discuss results for the slowest stretching speed simulations as good representatives of all the SMD simulations for this polarized desmosomal system.

**FIGURE 5.**
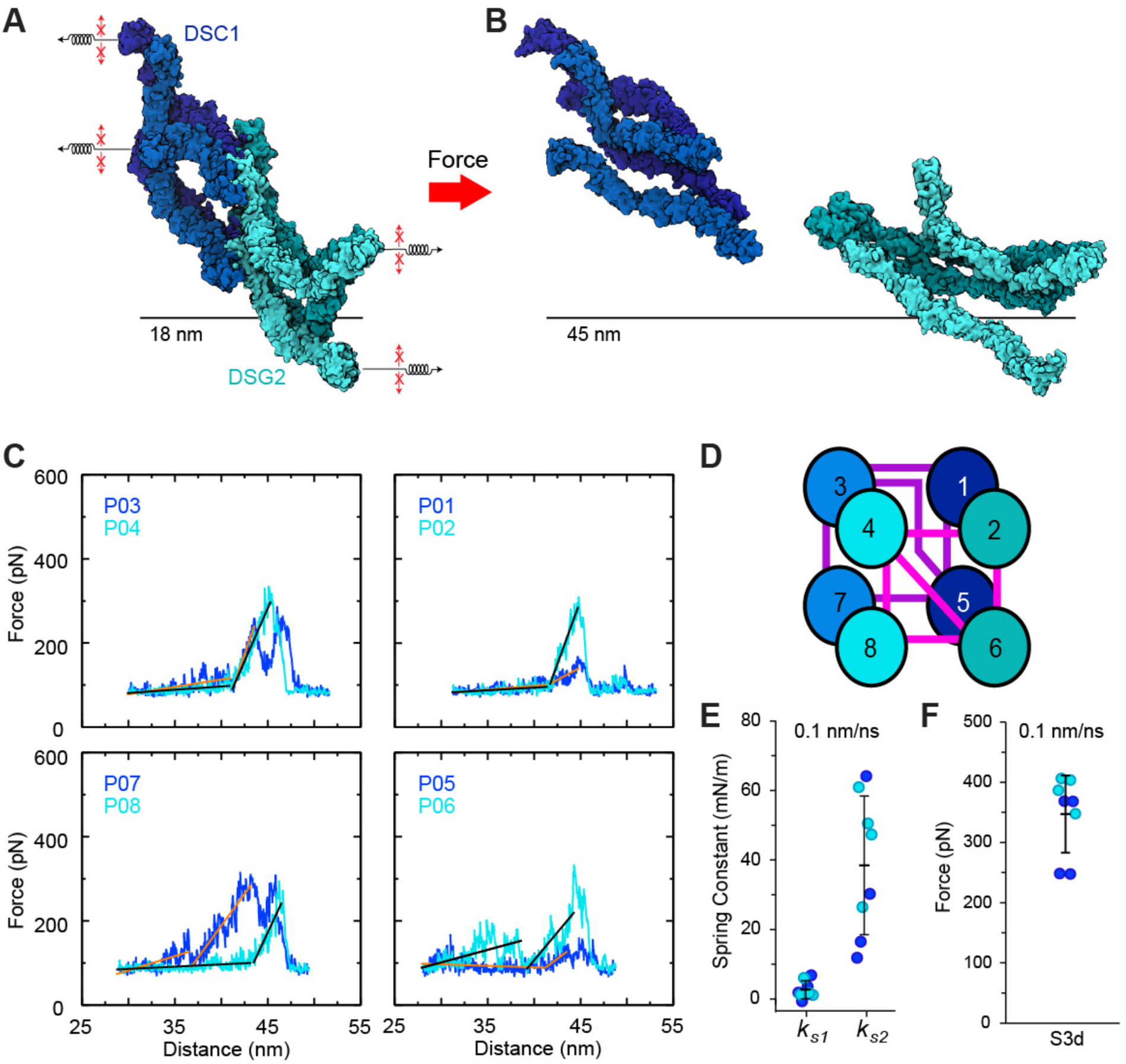
Forced unbinding of a polarized DSG2-DSC1 junction. (*A*) Starting state of the DSG2-DSC1 polarized junction. Springs with arrows indicate the stretching direction. Small red arrows indicate constraints in the plane applied to mimic cytoskeleton attachment. Black bar is the distance between hypothetical cellular planes. (*B*) A ruptured state of the DSG2-DSC1 junction (simulation S3d at 0.1 nm/ns; Table 1). (*C*) Force versus end-to-end distance plots for constant-velocity stretching of individual DSG2-DSC1 *trans* dimer pairs within the junction (0.1 nm/ns, simulation S3d). Bright blue lines are for DSG2 monomers on the right side of the junction in (*A*) and (*B*) or the top side of the junction in (*D*); blue lines are for DSC1 monomers on the left side in (*A*) and (*B*) or the bottom side of the junction in (*D*); black and orange lines are linear fits used to determine elasticity for DSG2 and DSC1, respectively. Plots are arranged to reflect the position of DSG2-DSC1 *trans* dimers within the junction, as labeled in (*D*) (P01 –P08). (*D*) Schematic of the DSG2-DSC1 junction (bright blue circles are DSG2; blue circles are DSC1). Positions of DSG2 and DSC1 monomers are labeled P01 – P08. Darker circles represent monomers on the backside of the system in (*A*) and (*B*). *Trans* interactions are represented by an overlap of circles. Lines indicate *cis* interactions: pink are for *cis* interactions among top layer monomers while purple are for *cis* interactions among bottom layer monomers. (*E*) Spring constants for phase 1 (*k_s_*_1_) and phase 2 (*k_s_*_2_) from simulation S3d. Values for individual monomers indicated as circles and colored according to type. Average and standard deviation shown as black bars. (*F*) Peak force required to rupture *trans* dimers in the DSG2-DSC1 junction. Circles and bars as in (*E*).

Similar to the 24-CDH1 junction, the polarized desmosomal junction exhibited a two-phased elastic response when stretched at a constant velocity of 0.1 nm/ns (simulation S3d; Fig. 5, *A* and *B*; Video S7). In the first phase, the DSG2-DSC1 *trans* dimers unbent at small forces (∼ 50 to 175 pN) over extensions of ∼ 5 to 10 nm with an associated soft spring constant of *k_s1_* = 2.7 ± 2.6 mN/m (simulation S3d; averages computed from slopes of force *versus* end-to-end distance curves on each side of the junction for all *trans* dimers including both DSG2 and DSC1 monomers; Fig. 5 *C* and *E*). After this initial soft extension mediated by unbending, the DSG2-DSC1 *trans* dimers became stiffer before rupturing, with a spring constant of *k_s2_* = 38.4 ± 20.0 mN/m. The spring constants associated with the first and second phases of extension for DSG2 and DSC1 monomers were not significantly different between the two types of proteins (*k_s1-DSG_* = 2.5 ± 2.3 mN/m and *k_s1-DSC_* = 2.9 ± 3.2 mN/m; *k_s2-DSG_* = 46.3 ± 14.5 mN/m and *k_s2-DSC_* = 30.6 ± 23.6 mN/m). Unlike the CDH1 *trans* dimers that were stiffer (larger *k_s2_*) within the 24-CDH1 adherens junction when compared to single isolated CDH1 *trans* dimers, the average spring constant for the second phase in extension in the desmosomal DSG2-DSC1 polarized junction was smaller than when computing it for individual isolated DSG2-DSC1 *trans* dimers (*k_s2_* ∼ 54.5 mN/m) at the same stretching speed, indicating that *cis* contacts did not stiffen but rather facilitated a more elastic mechanical response to force for the polarized desmosomal junction.

After unbending and stretching, unbinding for each of the *trans* dimers proceeded without unfolding when swapped Trp^2^ residues dislodged at the maximum force peak of *F*_p_ ∼ 347.1 ± 64.2 pN for simulation S3d at 0.1 nm/ns (average computed over forces monitored across all *trans* dimers on each side of the junction; Fig. 5 *F*). An exception occurred for the monomer at P03, which had its Trp^2^ residue already out of the pocket of its binding partner at P04 during equilibration. Interestingly, the unbinding force for the *trans* dimer pair at P03-P04 (∼299 pN) was still comparable to those of other dimers in the junction, suggesting that EC1-EC1 contacts such as those described above can help provide a robust response to force. As with CDH1 monomers within the 24-CDH1 adherens junction, disulfide bonds at the EC5 C-termini of DSG2 and DSC1 prevented unfolding, and rupture of individual *cis* and *trans* interactions manifested in various force peaks for the heterophilic *trans* dimers within the junction.

We monitored BSA for *trans* and *cis* interactions to distinguish and correlate their rupture to force peaks (simulation S3d; Fig. S6). For instance, when the end-to-end distance was ∼ 42 nm for the *trans* dimer at P05-P06, a partial break in the *cis* interaction between DSC1 EC1 at P05 and DSC1 EC2-EC3 at P07 correlated with a drop in the force applied to DSG2 at P06 (Fig. 5 *C*, lower right panel) and with a BSA drop from approximately 1000 Å^2^ to 300 Å^2^ between the monomers at P05 and P07 (Fig. S6, lower right panel, light orange). After the initial *cis* interaction between DSC1 monomers at P05 and P07 broke, a new stable *cis* contact was established between the same monomers with a different interface that contributed to an increase in BSA to ∼ 600 Å^2^ (Fig. S6, lower right panel, light orange). These *cis* interactions that were maintained throughout the unbinding trajectory resulted in a low rupture force for P05. Similarly, the successive rupture of *cis* contacts between DSG2 at P06 and DSG2 at P02 correlate with two force peaks for DSG2 at P06 prior to *trans* rupture with DSC1 at P05 (Fig. S6 *A*, lower right panel, red). Similar effects were observed throughout the lattice, with the breaking and formation of *cis* interactions modulating the force profile for individual monomers, resulting in complex force profiles with multiple force peaks for the DSG2-DSC1 *trans* dimers within the desmosomal junction. Regardless, we still observed a two-phased response with soft unbending of monomers followed by stiffening of their response to force before unbinding without unfolding upon dislodging of Trp^2^ residues.

#### Stiffer unbending and more resistant unbinding in a crisscross desmosome during tensile stretching

The second model representing a minimalistic desmosome junction also has the entire ectodomains of four DSG2 monomers on one side forming *trans* dimers with four DSC1 monomers on the opposite side, but these are arranged in a crisscross fashion based on a three-dimensional cryo-ET map of an intact human epidermis desmosome (39) (see Materials and Methods; Fig. 6 and S2 *C*). During the equilibration all *trans* contacts remained stable. Additionally, a new *cis* contact was formed between DSG2 monomers at P08 and P06. After equilibration, SMD simulations were carried out as with the polarized desmosomal junction at stretching speeds of 10 nm/ns and 1 nm/ns (simulations S4b-c; Video S8). Here we discuss results for the slowest stretching speed simulations as good representatives of all the SMD simulations for this crisscross desmosomal system.

**FIGURE 6.**
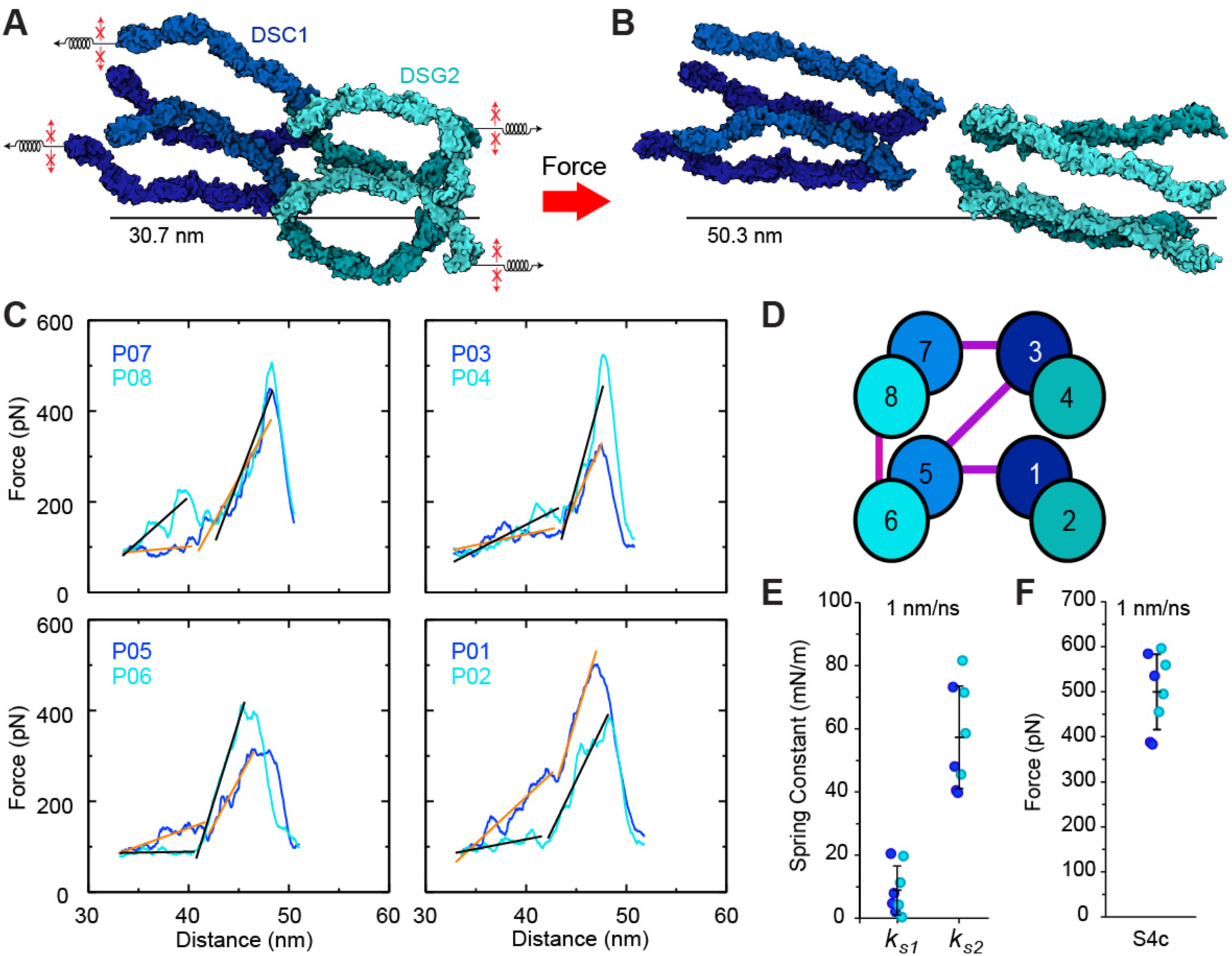
Forced unbinding of the crisscross DSG2-DSC1 junction. (*A*) Starting state of the DSG2-DSC1 crisscross junction. Springs with arrows indicate the stretching direction. Small red arrows indicate constraints in the plane applied to mimic cytoskeleton attachment. Black bar is the distance between hypothetical cellular planes. (*B*) A ruptured state of the DSG2-DSC1 junction (simulation S4c at 0.1 nm/ns; Table 1). (*C*) Force versus end-to-end distance plots for constant-velocity stretching of individual DSG2-DSC1 *trans* dimer pairs within the junction (1 nm/ns, simulation S4c). Bright blue lines are for DSG2 monomers on the right side of the junction in (*A*) and (*B*) or the top side of the junction in (*D*); blue lines are for DSC1 monomers on the left side in (*A*) and (*B*) or the bottom side of the junction in (*D*); black and orange lines are linear fits used to determine elasticity for DSG2 and DSC1, respectively. Plots are arranged to reflect the position of DSG2-DSC1 *trans* dimers within the junction, as labeled in (*D*) (P01 – P08). (*D*) Schematic of the DSG2-DSC1 junction (bright blue circles are DSG2; blue circles are DSC1). Positions of DSG2 and DSC1 monomers are labeled P01 – P08. Darker circles represent monomers on the backside of the system in (*A*) and (*B*). *Trans* interactions are represented by an overlap of circles. The pink line represents a *cis* interaction between DSG2 monomers and the purple lines indicate *cis* interactions among DSC1 molecules. (*E*) Spring constants for phase 1 (*k_s_*_1_) and phase 2 (*k_s_*_2_) from simulation S4c. Values for individual monomers indicated as circles and colored according to type. Average and standard deviation shown as black bars. (*F*) Peak force required to rupture *trans* dimers in the DSG2-DSC1 junction. Circles and bars as in (*E*).

Similar to the polarized desmosomal junction, the crisscross junction exhibited a two-phased elastic response, with a soft unbending phase preceding a stiffer response to force before unbinding without any unfolding of secondary structure. However, the degree of *cis* contacts is reduced in the crisscross junction compared to the polarized system (Fig. 6 *D*), and this difference was manifested in the elastic response to force. The spring constants associated with first and second phase of the crisscross junction were *k_s1_* ∼ 8.8 ± 7.7 mN/m and *k_s2_* ∼ 57.3 ± 16.3 mN/m, respectively (Fig. 6 *C* and *E*), both of which are stiffer than the spring constants for the polarized junction when stretched at the same speed (*k_s1_* ∼ 3.5 ± 2.9 mN/m and *k_s2_* ∼ 44.3 ± 31.7 mN/m at 1 nm/ns). This difference in elasticity can be attributed to the reduction in *cis* contacts of the crisscross system, which results in a greater proportion of force applied to the *trans* interactions as opposed to being distributed through the lattice via *cis* contacts. This structural difference also results in larger force peaks during unbinding, with an average of *F*_p_ ∼ 499.7 ± 83.9 pN for simulation S4c (average computed over forces monitored across all *trans* dimers on each side of the junction; Fig. 6 *C* and *F*), which was larger than the average force monitored for the polarized complex at the same speed (*F*_p_ ∼ 373.4 ± 75.9 pN, simulation S3c). Based on our simulations, a junction formed by the crisscross configuration would be stiffer and stronger, while a junction formed in a polarized configuration would be more elastic and less resistant to force (Fig. 8). These differences may help further elucidate what type of architecture is observed *in vivo* (42).

### Brittle Mechanical Response of a Clustered PCDH Junction

Clustered PCDH proteins are expressed in the brain, where neuronal tissue is exposed to different types of mechanical stimuli than those experienced by cardiac and epithelial tissues (66–73, 76). Simulations of clustered PCDH *trans* dimers reported in a companion manuscript show that these complexes lack the soft unbending phase found for classical cadherins and that these unbound at larger forces, suggesting a brittle response that may underlie their functional role in neuronal self-avoidance and non-self discrimination. However, recent cryo-ET data of clustered PCDHs in liposome junctions and an X-ray crystallographic structure of the full-length ectodomain of PCDHγB4 (99) suggest that the clustered PCDH *trans* dimers form a lattice with an architecture that might provide some resilience, as EC6-mediated *cis* dimers emerge from the membrane forming “V” shaped units that could close their aperture upon application of tensile forces. To test the elasticity of clustered PCDHs within junctions, we built a model with eight PCDHγB4 monomers forming three *trans* dimers with four *cis* contacts. This PCDHγB4 junction was equilibrated for 20 ns with constraints applied to C-terminal C_α_ atoms (simulation S5a). The large anti-parallel *trans* interface and the smaller *cis* interactions were maintained during equilibration, which was followed by SMD simulations in which C-termini from monomers in opposite sides of the junction were moved in opposite directions to induce tensile stretching and unbinding. We performed constant velocity SMD on this system at three different speeds; 10, 1, and 0.1 nm/ns (simulations S5b-d). During these simulations, additional harmonic constraints were applied to the C-terminal C_α_ atoms in the plane perpendicular to the stretching axis to mimic attachment to cytoskeletal elements. At the fastest stretching speed of 10 nm/ns, individual PCDHγB4 monomers unfolded while unbinding from their *trans* binding partners, however at slower stretching speeds of 1 nm/ns and 0.1 nm/ns we observed unbinding without unfolding. We discuss results for the slowest stretching speed simulation as a good representative of the slower SMD simulations for this clustered PCDH junction.

The mechanical response of the PCDHγB4 junction stretched at a constant velocity of 0.1 nm/ns (simulation S5d; Fig. 7 *A – F*, Videos S9 and S10) was markedly different than the response monitored for the other simulated classical cadherin junctions. There was a noticeable lack of a two-phased elastic response, with monomers displaying a short (< 2 nm) initial “soft” response to tensile forces with an associated spring constant of *k_s1_* = 35.2 ± 12.3 mN/m (Fig. 7 *E* and *H*), comparable to the stiffer second phase observed for the 24-CDH1 and desmosomal junctions in simulations at their slowest stretching speeds. The predominant second phase, characterized by a spring constant of *k_s2_* = 148.5 ± 32.4 mN/m over an extension of ∼ 4 nm (Fig. 7 *E* and *H*; Video S9), was associated with the closure of the “V” shaped *cis* dimers and subsequent stretching of the large and rigid *trans* interfaces, followed by unbinding at forces averaging *F*_p_ ∼ 607.9 ± 146.3 pN (Fig. 7, *A-C* and *I*), almost double the average force required to rupture the classical cadherin junctions in our simulations stretched at the same (polarized desmosome) or faster speeds (adherens junction). This suggests that unbinding forces are in general higher for the PCDHγB4 junction compared to classical cadherin junctions, since peak forces required to rupture complexes in general decrease with decreasing stretching speeds (135).

**FIGURE 7.**
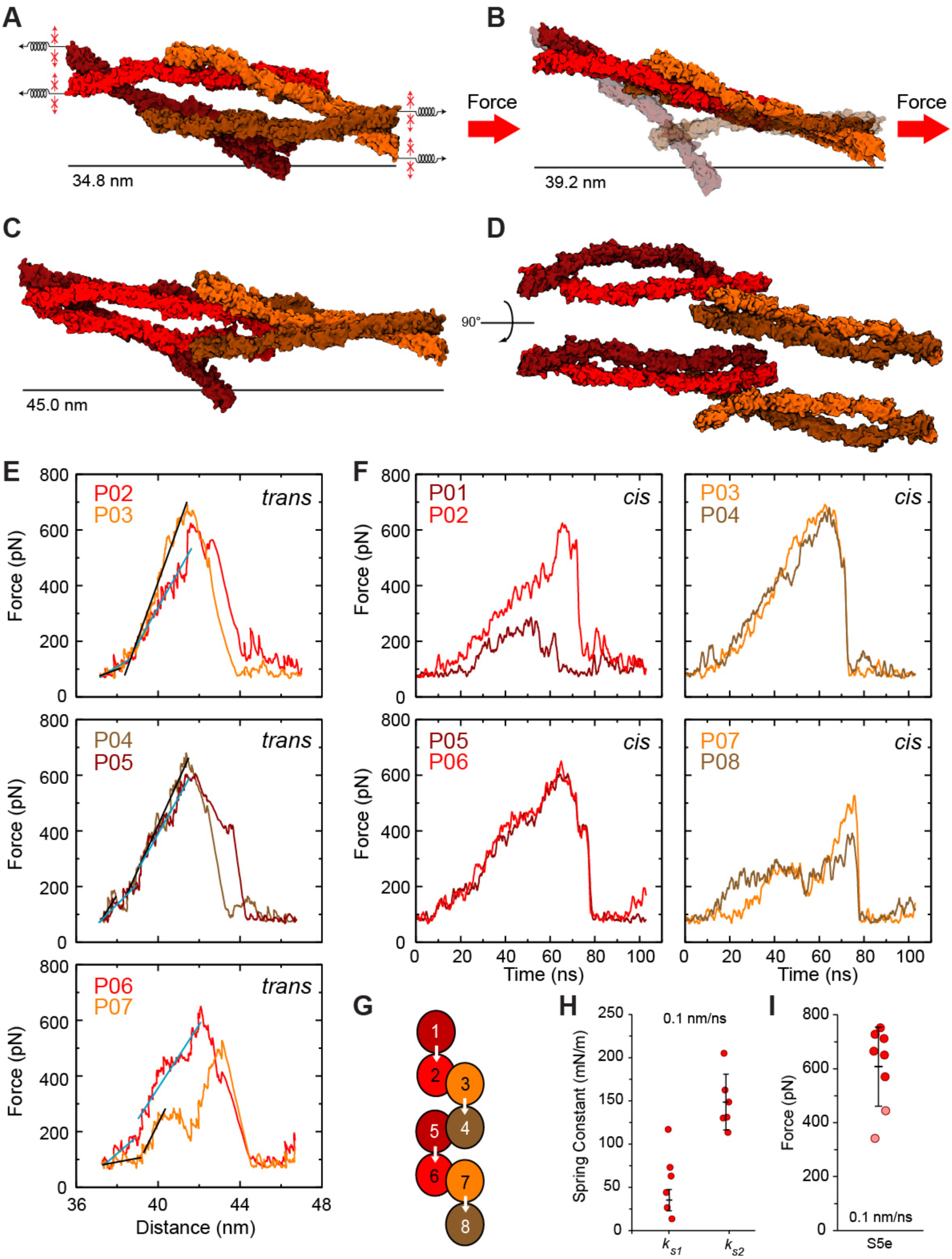
Force unbinding of PCDHγB4 junction. (*A*) Side view of the starting state of the PCDHγB4 junction. Small red arrows indicate constraints in the plane applied to mimic cytoskeleton attachment. Black bar is the distance between hypothetical cellular planes. (*B*) Side view of the stretched state of the PCDHγB4 junction highlighting the closure of the aperture facilitated by *cis* interactions (simulation S5d at 0.1 nm/ns; Table 1). Monomers not involved in *trans* interactions are transparent. (*C*) Final state of the PCDHγB4 system after stretching at 0.1 nm/ns (simulation S5d). (*D*) A 90° rotate view of (*C*). (*E*) Force versus end-to-end distance plots for constant velocity stretching of individual PCDHγB4 *trans* dimer pairs within the junction (0.1 nm/ns, simulation S5d). Bright and light colors for force curves of PCDHγB4 monomers alternate according to their location in the lattice as indicated. Cyan and black lines are linear fits used to determine elasticity for PCDHγB4 monomers on the left or right side in (*A*-*D*), respectively. Plots are arranged to reflect the position of PCDHγB4 *trans* dimers within the junction, as labeled in (*G*) (P01 – P08). (*F*) Force versus time plots for constant velocity stretching of individual PCDHγB4 *trans* dimer pairs grouped by *cis*-interacting pairs. Data labeled as in (*E*). (*G*) Schematic of the PCDHγB4 junction. Positions of monomers are labeled P01 – P08. *Trans* interactions are represented by an overlap of circles. White arrows indicate *cis* interactions. Arrows show *cis* interaction directionality with their base representing EC5 of the originating monomer and the arrowhead representing the binding surface of EC6 on another monomer. (*H*) Spring constants for phase 1 (*k_s_*_1_) and phase 2 (*k_s_*_2_) from simulation S5d. Values for individual monomers indicated as circles. Pale circles are for monomers not involved in *trans* interactions. Average and standard deviation shown as black bars. (*I*) Peak force required to rupture the *trans* dimers in the PCDHγB4 junction. Circles and bars as in (*H*). Pale red circles represent values obtained for PCDHγB4 monomers not involved in *trans* interactions.

After the initial rupture of the *trans* interaction, monomers slipped past each other with very little resistance, except for small force peaks associated with transient *trans* intermediates formed by EC2-EC2 contacts between monomers at P06 and P07, as well as monomers at P04 and P05 (Fig. 7 *D* and *F*, lower panels; Fig. S7 *A*, middle and lower panels; Video S10). The *cis* contacts mediated by EC6 remained throughout the trajectory, acting as hinges that triggered the re-opening of the V-shaped *cis* dimers after complete unbinding. Interestingly, in addition to EC6-EC6 *cis* interfaces that were present from the beginning, two pairs of monomers, at positions P03 and P04 as well as P05 and P06, formed new *cis* interfaces involving repeats EC1 and EC2 as indicated by the increase in their BSA during stretching (Fig. S7 *B*). Overall, the response to tensile forces of PCDHγB4 monomers within the junction recapitulated what we observed for the response of other clustered PCDH *trans* dimers (PCDHα7, PCDHβ6, and PCDHγB3), suggesting that these form brittle junctions.

## DISCUSSION

The dynamics and mechanics of cadherin junctions *in situ* can be difficult to study experimentally at high resolution, yet these cell-cell junctions are at the core of tissue function in various mechanical processes, including normal stretching and shearing during morphogenesis and cardiovascular activity, as well as in more extreme circumstances such as responses to external abrasions and to mechanical insults. Modeling and simulation provide an attractive alternative approach to studying the structure, dynamics, and mechanics of multi-protein arrays (113, 136–138), including of cadherins complexes. Indeed, some classical cadherin-based junctions have been simulated previously and were modeled as either coarse-grained (139), rigid-body (56, 140), or all-atom (42) systems including only the adhesive cadherin ectodomains. These simulations showed that there might be a change in cadherin inter-monomer spacing when cellular planes of adherens junctions are pulled apart (139), that the formation of an adherens junction relies on the positive cooperativity of *cis* and *trans* interactions (56, 140), and that the initial arrangement of DSG2-DSC2 *trans* dimers within a desmosomal junction can alter its overall mechanical response, an effect that can be used to determine viable junction architectures (42). Our MD simulations of cadherin ectodomains, add an all-atom view on the mechanics of CDH1-based adherens junctions, report on the predicted strength and unbinding pathways for DSG2-DSC1 *trans* cadherin dimers within two arrangement types of desmosomal junctions, and test the unexplored mechanics of a non-classical clustered PCDH junction formed by PCDHγB4. These simulations provide key insights on the mechanics of each multi-cadherin system, while the combined set allows us to compare their responses in the context of their different functions (Fig. 8).

**FIGURE 8.**
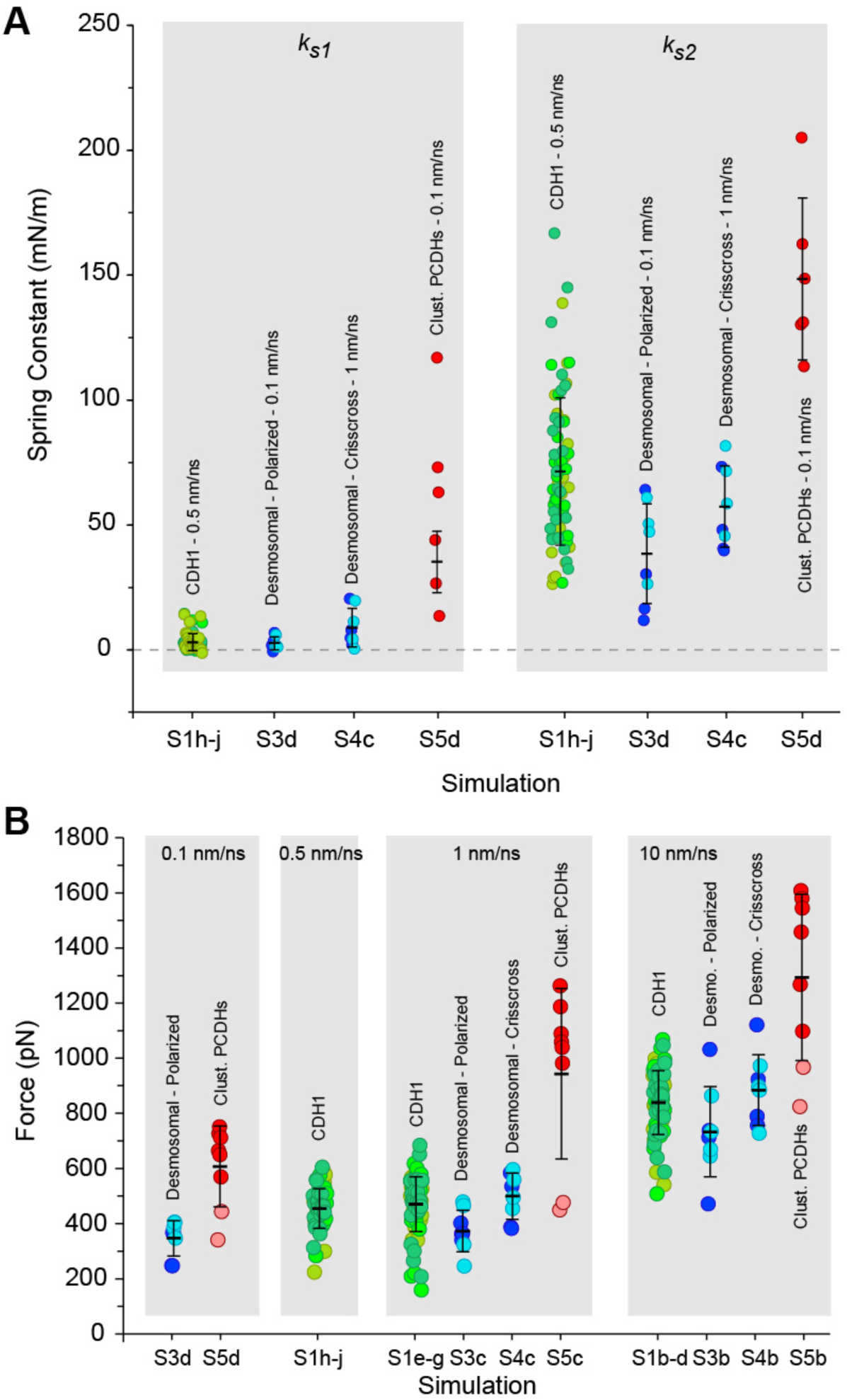
Predicted elasticity and peak rupture forces for cadherins within junctions. (*A*) Summary of values obtained for the spring constants for the soft (*k_s1_*) and stiff (*k_s2_*) phases. (*B*) Summary of peak unbinding forces grouped by stretching speeds. Pale red circles represent values obtained for PCDHγB4 monomers not involved in *trans* interactions.

In a companion paper we show that the simulated response of individual classical cadherin *trans* dimers to tensile stretching is characterized by two phases with soft straightening of ectodomains over ∼ 10 nm extensions followed by stiffening that leads to unbinding without unfolding. On the other hand, individual clustered PCDH *trans* dimers were brittle and lacked the initial soft straightening phase. Here we observe a similar trend when these *trans* dimers are embedded within a junction, forming *cis* contacts with other cadherin monomers. Interestingly, *cis* contacts in classical cadherins, mainly mediated by EC1 and EC2 repeats, tune the monomer’s responses to tensile stretching, adding a layer of complexity to force profiles that often have various force peaks as some *trans* dimers pick up the slack of neighbors through *cis* interactions. The behavior of classical cadherin *trans* dimers within a junction is more heterogeneous than the behavior of isolated complexes, and seems to be dependent on their junction location, with those at the periphery often having different force profiles than those at the center. Nevertheless, most force profiles still displayed two phases with soft stretching over ∼ 5 to ∼ 10 nm. In contrast, the clustered PCDHγB4 *trans* dimers with EC6-mediated *cis* contacts displayed a more homogeneous brittle response within the junction. Although the V-shaped clustered PCDHγB4 *cis* dimers flattened upon tensile stretching, the *cis* contacts did not break. There was little extension upon tensile stretching of *trans* dimers within the PCDH junction before rupture, and forces needed to trigger *trans* unbinding were larger than what we observed for classical cadherin junctions (Fig. 8). As with the classical cadherin *trans* dimers, subsequent unbinding occurred without unfolding at the slowest stretching speed. These results confirm that classical cadherins tend to be mechanically resilient while clustered PCDH ectodomains tend to be brittle, even within junctions.

Our simulations also show that *cis* interactions within classical cadherin junctions are more dynamic than *trans* interactions during tensile stretching and relaxation. During stretching, *cis* interactions ruptured, partially released, and re-formed (Figs. S5 and S6), quickly recovering during relaxation after stretching, even during the relaxation of the *trans*-ruptured stretched system (Fig. 3 *E*). In contrast, *trans* interactions were the last to break and, for the systems tested, did not fully recover after rupture in the short timescale of our simulations. Rapid recovery of stretched but not of *trans* ruptured adherens junctions suggest that these may act as molecular shock absorbers in short time-scales, although the extent of their reversible extension might be limited, with about 0.1 % of reversible tissue extensibility contributed from ectodomains alone (∼ 10 nm for a junction between two cells, ∼ 10 μm for 1000 cells lined up without taking into account any compression of the complex).

While there is still some uncertainty on how the ectodomains of desmosomal and clustered protocadherins form junctions *in vivo* (38, 39, 42, 99), the architecture of adherens junction has been more thoroughly studied (40) and thus a model of the 16-CDH1 junction was further tested in simulations that induced shearing rather than tensile stretching. These simulations revealed that our minimalistic adherens junction model has a preferential shearing direction in which separation and rupture require less work than when shearing forces are applied in the opposite direction (Fig. 4). Large adherens junctions are composed of many patches of smaller CDH1-based junctions (44) that might align in various orientations to offset this potential mechanical weakness.

The two desmosomal junction architectures tested in our simulations exhibited divergent mechanical properties, with the crisscross configuration behaving more similar to isolated desmosomal *trans* dimers with higher spring constants in both unbending phases, and higher rupture forces compared to the polarized system (Fig. 8). Desmosomes are known to exist in two states characterized by their susceptibility to Ca^2+^ chelators (141). While Ca^2+^-dependent desmosomes might be weak, Ca^2+^-independent “hyperadhesive” desmosomal junctions might withstand strong forces in the epidermis, the throat, tongue, liver, and cardiomyocytes *in vivo* (141). Given that hyperadhesiveness seems to rely on external factors and maturation timescales that exceed the short time scale of atomistic MD simulations (110, 111), it is unlikely that our simulations alone can provide insights into the molecular architecture of the hyperadhesive state. Most likely we are probing the elastic and mechanical response of newly formed desmosomes (142) or desmosomes present in tissue weakened by previous nearby trauma (143) and implicated in the process of wound healing. The dynamic nature of the *cis* interactions observed in the slowest forced unbinding of the polarized desmosomal junction, with *cis* contacts breaking and reforming for both DSG2 and DSC1 pairs (Fig. S6), may help facilitate desmosomal rearrangements needed to establish a hyperadhesive state after the tissue has sustained a wound, but other aspects, including changes in molecular composition (142), might determine desmosome maturation into Ca^2+^-independent states.

The predicted mechanical strength of the PCDHγB4 *trans* dimers, isolated and in junctions, is intriguing given that the clustered PCDHs mainly play a role in neuronal recognition (82, 83), while the adherens junction and the desmosome maintain epithelial cell-cell adhesion and must withstand significant mechanical stress. The γ-PCDHs, of which PCHDγB4 is a member, might play a role in adhesion at the synapse (98, 144), but it is difficult to say if the clustered PCDHs are under or respond to any mechanical strain *in vivo*. Perhaps this brittle response is appropriate for more subtle types of mechanical stimuli that clustered PCDHs may experience in neurons during development, normal function, and in brain injury (66–73, 76).

The distinct mechanical behaviors of classical cadherins and clustered protocadherins predicted by our simulations might also have implications for assembly and differential survival of the junctions they form in various types of tissues. Tissues and individual cell membranes do experience oscillatory motions (145–149), with amplitudes of up to 3 nm and characteristic timescales that can be slow (hundreds of milliseconds) (149) or fast (hundreds of microseconds) (145). These observations suggest that the junctions we simulated would be subject to periodic stretching and compression. The two-phased response exhibited both by the CDH1 and desmosomal junctions as well as buffering of mechanical perturbation by *cis* interactions seen in our simulations offer a molecular explanation of how cell-cell junctions may withstand this oscillatory perturbation observed *in vivo*. In mammalian neuronal systems, propagation of action potentials are accompanied by cellular deformations of up to ∼ 3 nm at sub-millisecond timescales (70–75), which could induce rupture of the brittle clustered PCDH junctions. These admittedly speculative scenarios suggest that mechanical motions triggered by cellular activity may control cell-cell adhesion, not only by selecting for adhesive junctions that can withstand a given mechanical stimulus, but also perhaps by facilitating fluctuation driven assembly of junctions through a ratchet-like mechanism (150). Our simulations suggest that, in addition to quantity and affinity (151), the molecular mechanics of adhesion and the activity of cells might be relevant for cell sorting and connectivity (152, 153).

From an evolutionary perspective, it is interesting to speculate that ancestral cadherins with long and likely flexible ectodomains (21, 133, 154) might have been replaced by modern cadherins with shorter but mechanically equivalent ectodomains. The evolutionary transition from an aquatic based lifestyle to land was accompanied by an increase in desmosomal protein diversity (155), suggesting that desmosome expansion was required for further development of a more robust junction to resist perturbations associated with a land-based lifestyle. Yet whether mechanical force has played a role in shaping the molecular evolution of various cadherin junctions, including desmosomes, remains to be explored.

Overall, our work provides an atomistic exploration of how three essential cadherin-based cell-cell junctions respond to force as aggregate units, and how *cis* interactions modulate the properties of these junctions at all stages of the unbinding trajectory. While simulated timescales are short (hundreds of nanoseconds), the simulated conditions used are equivalent to those experienced by tissues exposed to blunt trauma (126, 127), and we expect that our quantitative predictions from stretching simulations at fast speeds will provide upper bounds for elasticity and unbinding forces, with qualitative predictions holding even at near equilibrium conditions (103, 104, 110, 156–158). Future modeling efforts should focus on taking into account the effect of Ca^2+^, glycosylation, membrane, and cytoplasmic partners on the tensile and shearing mechanics of junctions that might not only include mixtures of cadherin proteins, such as classical and clustered or delta-protocadherins (159–161), but also complexes with neuroligins, nectins, and other proteins partners (97, 162–165).

## SUPPORTING INFORMATION

FIGURES S1-S7 and VIDEOS S1 to S10 are available at are available at XXX

## AUTHOR CONTRIBUTIONS

BLN and RAS prepared and simulated CDH1 systems. CN prepared and simulated systems with desmosomal cadherins. SW and MS prepared the clustered PCDH system, which was simulated by SW. BLN analyzed data with input from all co-authors. MS trained co-authors and supervised research. BLN, CN, SW, and MS designed research and wrote and edited the manuscript.

## ACKNOWLEDGEMENTS

This work was supported by the Ohio State University and by the Human Frontier Science Program (RGP0056/2018). Simulations were performed using the NCSA-Blue Waters (GLCPC), TACC-Stampede, PSC-Bridges (XSEDE MCB140226), OSC-Owens, and OSC-Pitzer (PAS1037 and PAA0217) supercomputers. B. L. N. was supported by an OSU/NIH cellular, molecular biochemical sciences program training grant fellowship (T32GM086252) and by an OSU presidential fellowship. C. N. was supported by an OSU/NIH molecular biophysics training grant (TG32GM118291). R. A.-S. was a Pelotonia fellow. M. S. was an Alfred P. Sloan fellow (FR-2015-6794).

