## Supplement for "Collective Mechanical Responses of Cadherin-Based Adhesive Junctions as Predicted by Simulations"

July 2021

**Video S1**. Side view of a 24-CDH1 junction under tensile forces with straightening and unbinding. Monomers were stretched at 0.5 nm/ns (simulation S1h in Table 1, 0 – 49.0 ns). Initial straightening was followed by rupture of *cis* interactions between neighboring monomers at both sides of the junction. Towards the end of the simulation *trans* interactions are ruptured without unfolding. The unbound CDH1 molecules begin the re-bend. Proteins are depicted in a rough surface representation (greens). Water molecules and other atoms are not shown for clarity.

**Video S2**. Front view of a 24-CDH1 junction under tensile forces with straightening and unbinding. Simulation trajectory shown in Video S1 from a different viewpoint (simulation S1h in Table 1, 0 – 49.0 ns). System is displayed as in Video S1.

**Video S3**. Relaxation of a CDH1 24-molecule system after 21.1 ns of stretching at 1 nm/ns (simulation S1k). The initial starting state maintained all *trans* interactions and had five *cis* interactions remaining. During the trajectory the straightened CDH1 monomers begin to quickly re-bend and some *cis* interactions re-form. System is displayed as in Video S1.

**Video S4**. Relaxation of a CDH1 24-molecule system after 23.1 ns of stretching at 1 nm/ns (simulation S1l). The initial starting state had nine *trans* interactions and two *cis* interactions remaining. During the trajectory the individual CDH1 monomers began to quickly re-bend, one *trans* interaction re-formed without $Trp^2$ exchange while two additional *trans* interactions were lost, and some *cis* interactions re-formed. System is displayed as in Video S1.

**Video S5**. Shearing of a 16-CDH1 junction. Monomers were stretched at 0.5 nm/ns (simulation S2g in Table 1, 0 – 38.3 ns) in a direction that follows the natural tilt of the CDH1 *trans* dimers. Monomers straightened and *tran*s dimers ruptured. System is shown as in Movie S1.

**Video S6**. Shearing of a 16-CDH1 junction. Monomers were stretched at 0.5 nm/ns (simulation S2f in Table 1, 0 – 141.6 ns) in a direction that is against the natural tilt of the CDH1 *trans* dimers. Monomers are compressed first, with rupture of *cis* interactions as the system flips and monomers decompress. Monomers continued to be stretched until loss of *trans* interactions. System is shown as in Movie S1.

**Video S7**. Forced unbending and unbinding of DSG2-DSC1 *trans* dimers in a polarized desmosomal junction. Monomers were stretched at 0.1 nm/ns (simulation S3d in Table 1, 0 – 270.4 ns). Initial straightening was followed by rupture of *cis* and *trans* interactions without unfolding. Proteins are depicted in a rough surface representation (DSG2 – cyan; DSC1 – blue). Water molecules and other atoms are not shown for clarity.

**Video S8**. Forced unbending and unbinding of DSG2-DSC1 *trans* dimers in a crisscross junction. Monomers were stretched at 1 nm/ns (simulation S4c in Table 1, 0 – 19.8 ns). Initial straightening was followed by rupture of *cis* and *trans* interactions without unfolding. System is shown as in Video S7.

**Video S9**. Forced unbinding of PCDHγB4 *trans* dimers in a clustered PCDH junction. Monomers were stretched at 0.1 nm/ns (simulation S5d in Table 1, 0 – 103.4 ns). Flattening of the system is followed by rupture of *trans* interactions and the formation of new transient *trans* interactions as monomers slide past one another. The *cis* interactions are maintained throughout the course of the trajectory and cause the system to recover its V shape after the initial *trans* interactions are lost. Proteins are depicted in a rough surface representation (PCDHγB4 – reds, orange, brown). Water molecules and other atoms are not shown for clarity.

**Video S10**. Top view of Video S9 to highlight the nascent *trans* bond between EC2-EC2 of PCDHγB4 monomers at positions P06 and P07 (bottom). System displayed as in Video S9.

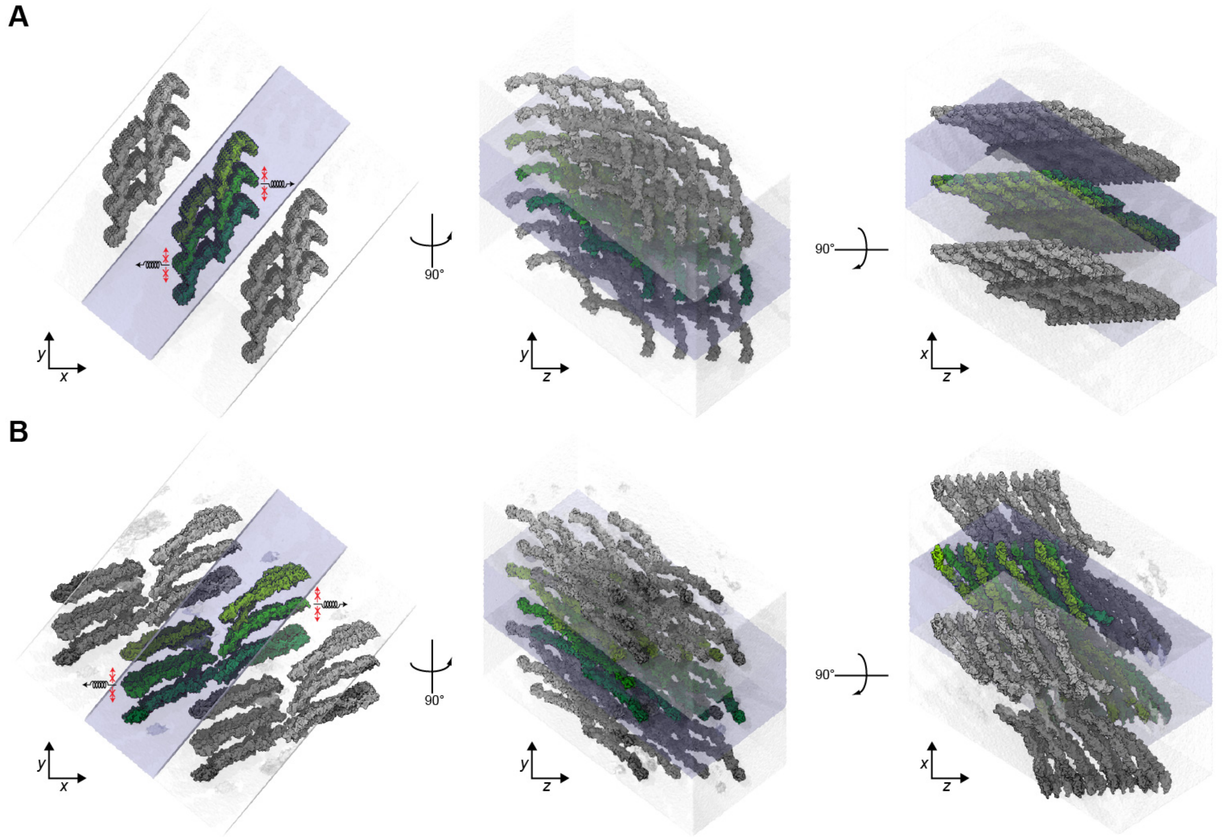


**FIGURE S1. System setup and periodic boundary conditions for the 24-CDH1 junction.** (*A*) The 24-CDH1 junction (green colors) is shown with two of its periodic images (greys) in three different orientations. Central water box is shown as a transparent blue surface, with adjacent periodic boxes in light gray. Springs with arrows (left panel) show the stretching direction along the *x* axis, red arrows indicate harmonic constraints applied in the plane of the hypothetical cell membrane to mimic attachment to the underlying actin cytoskeleton. (*B*) Snapshot of the 24-CDH1 junction at the end of simulation S1h show as in (*A*). The stretched CDH1 monomers cross the water box boundary to enter the space vacated by their periodic molecules. This arrangement allowed for systems to be simulated with smaller solvation boxes and stretching directions set to a primary axis.

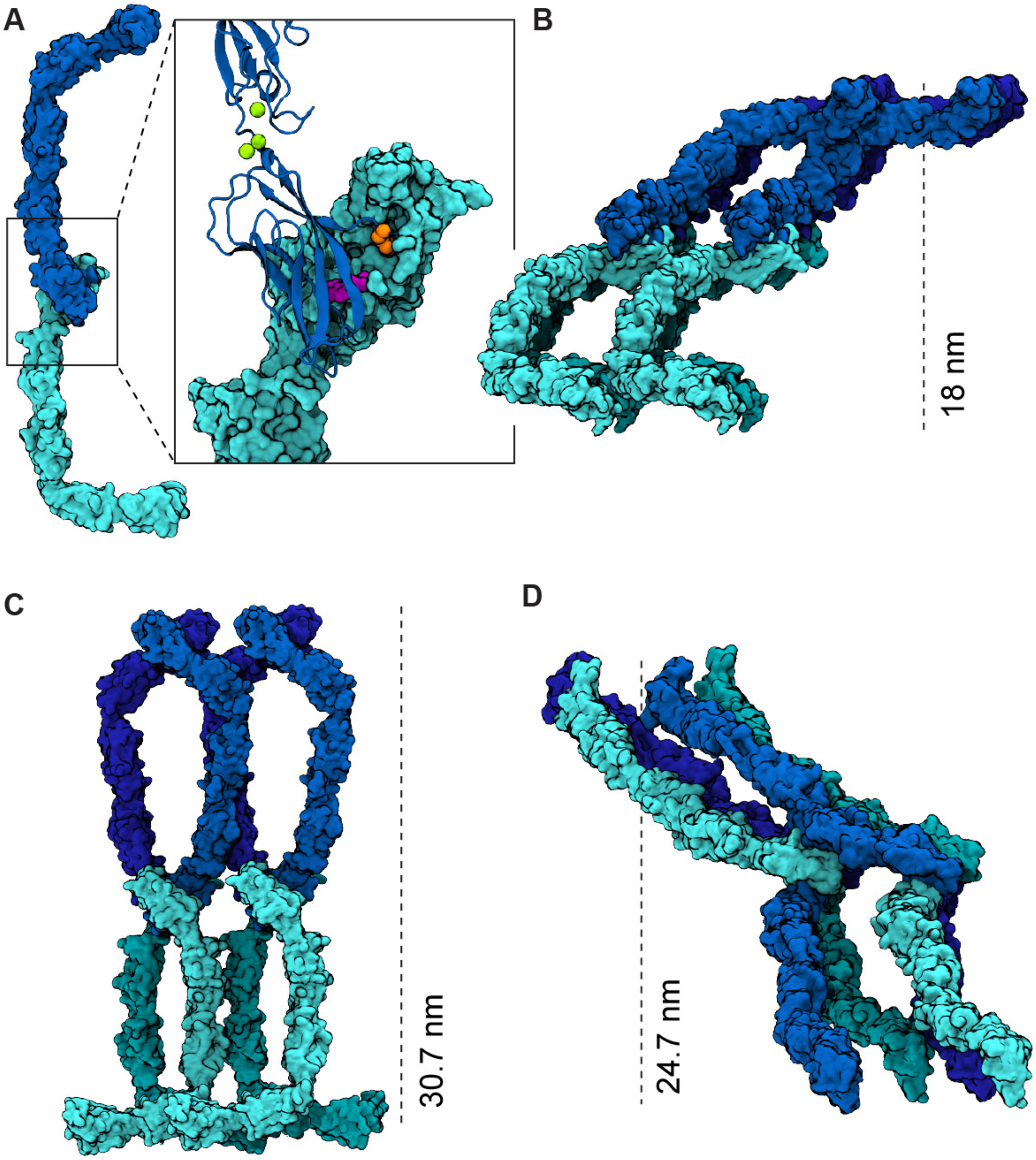


**FIGURE S2. Setup for the DSG2-DSC1 desmosomal junction.** (*A*) A DSG2-DSC1 *trans* dimer (DSG2 – cyan; DSC1 – blue) made by aligning the first six $C_\alpha$ atoms of the DSC1 monomer with the first six $C_\alpha$ atoms of the DSG2 monomer in the DSG2-DSG2 crystal homodimer structure (41). Inset shows the strand swap of $Trp^2$ from DSG2 into the hydrophobic binding pocket of DSC1 (DSG2 $Trp^2$ – purple; DSC1 $Trp^2$ – orange). $Ca^{2+}$ ions are shown in green, and DSC1 is shown in blue ribbon. (*B*) The polarized DSG2-DSC1 desmosomal junction. Distance between hypothetical cellular planes defined by C-termini is indicated by a dash line. (*C*) The crisscross DSG2-DSC1 desmosomal junction based on the arrangement proposed in (39) (*D*) The checkerboard DSG2-DSC1 desmosomal junction. This system was not simulated as serious clashes between monomer could not be easily resolved.

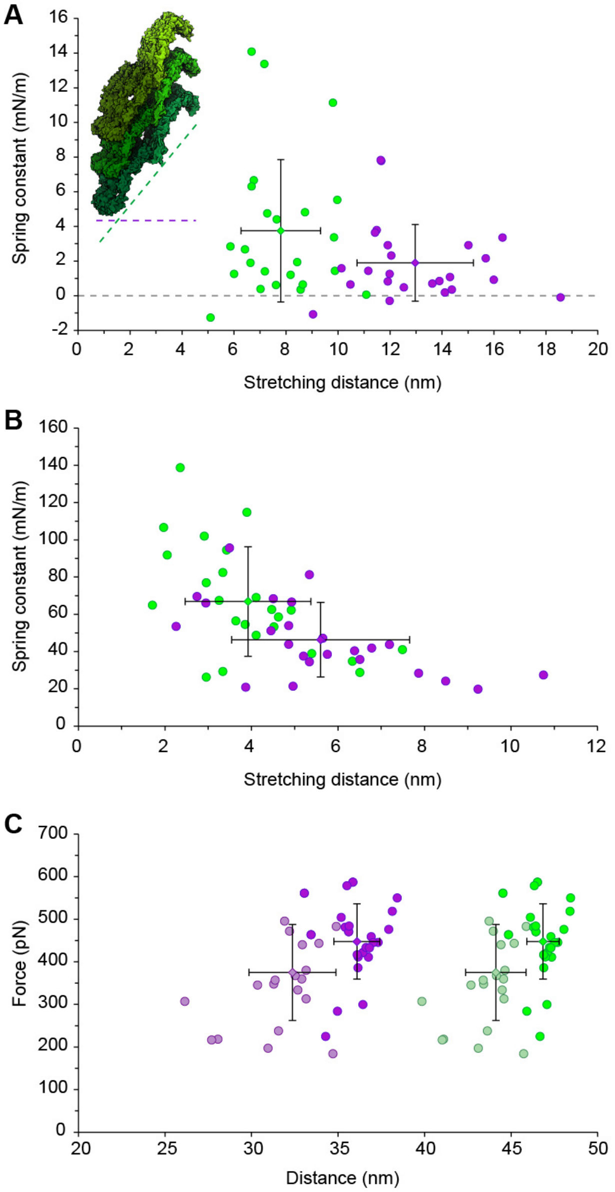


**FIGURE S3. Properties of 24-CDH1 junction as a function of end-to-end versus membrane-to-membrane distances.** (*A*) Plot of the first spring constants ($k_{s1}$) versus the distance of extension. The average spring constant is softer for membrane-to-membrane measurements ($k_{s1}$ = 1.9 ± 2.2 mN/m; extension = 13.0 ± 2.2 nm; simulation S1h; purple circles) compared to end-to-end measurements ($k_{s1}$ = 3.7 ± 4.1 mN/m; extension = 8.0 ± 1.5 nm; simulation S1h; green circles). Inset shows the 24-CDH1 junction system with dashed lines representing how the stretch distance was measured. (*B*) Plot of the second spring constant ($k_{s2}$) versus the distance of extension. The average spring constant is softer for membrane-to-membrane measurements ($k_{s2}$ = 46.3 ± 20.0 mN/m; extension = 5.6 ± 2.1 nm; simulation S1h) compared to end-to-end measurements ($k_{s2}$ = 66.9 ± 29.4 mN/m; extension = 3.9 ± 1.5 nm; simulation S1h). Colors are the same as in (*A*). (*C*) Peak force versus distance of stretch in the 24-CDH1 junction (simulation 1h). The maximum force peak (447.8 ± 88.3 pN) is shown versus end-to-end distance (46.8 ± 0.9 nm) and versus the membrane-to-membrane distance (36.1 ± 1.3 nm). The first force peak (375.2 ± 112.7 pN; pale circles) is shown versus end-to-end distance (at 36.1 ± 1.3 nm) and versus the membrane-to-membrane distance (at 32.4 ± 2.5 nm). Colors are the same as in (*A*).

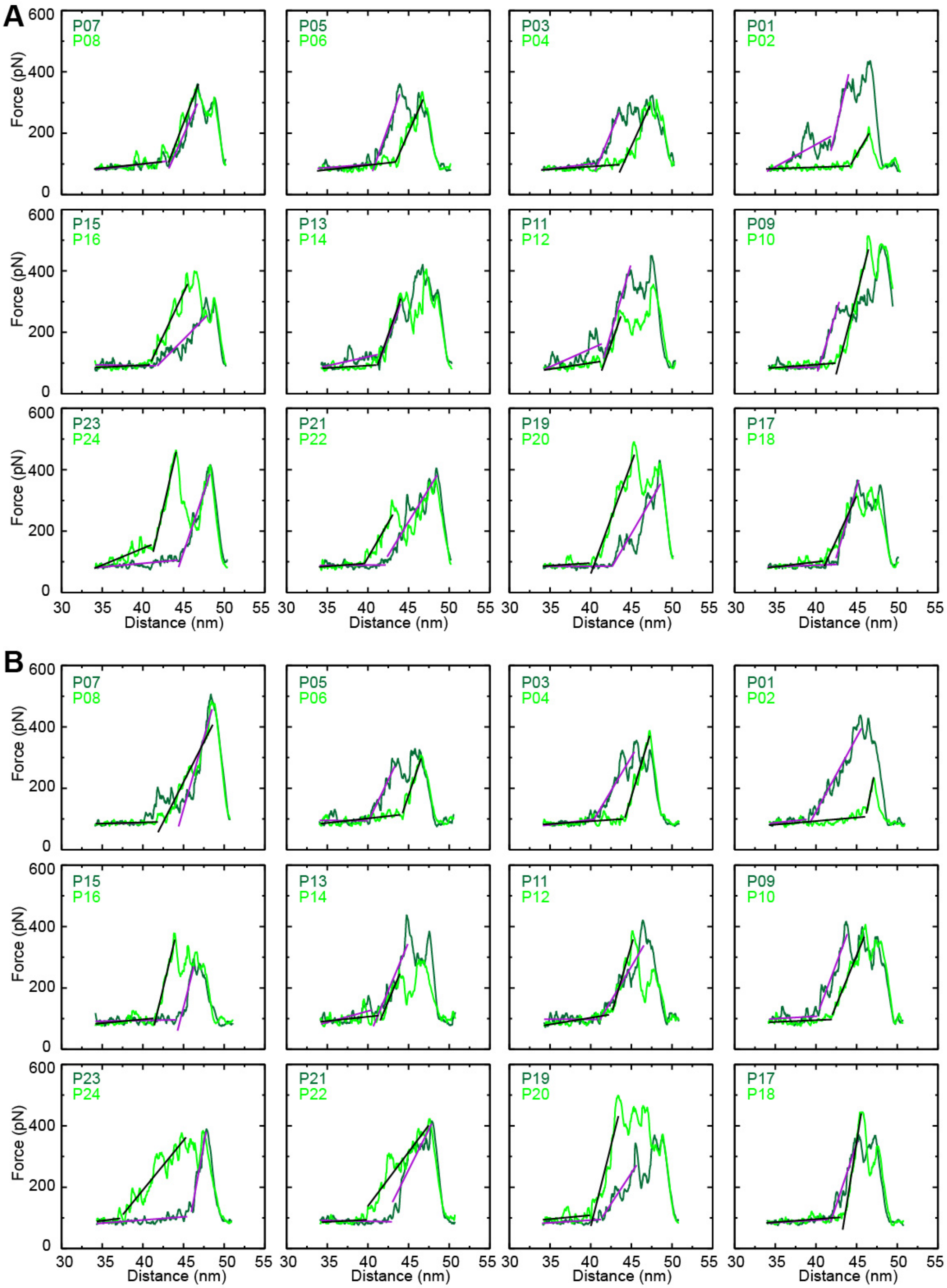


**FIGURE S4. Force profiles for unbinding of *trans* dimers in the 24-CDH1 adherens junction**. (*A*-*B*) Force versus end-to-end distance plots for constant-velocity stretching of individual CDH1 *trans* dimer pairs within the junction (0.5 nm/ns) shown as in Fig. 2 *C*. Data shown for simulations S1i (*A*) and S1j (*B*). Trends observed here and in Fig. 2 are robust despite difference in starting points for each simulation.

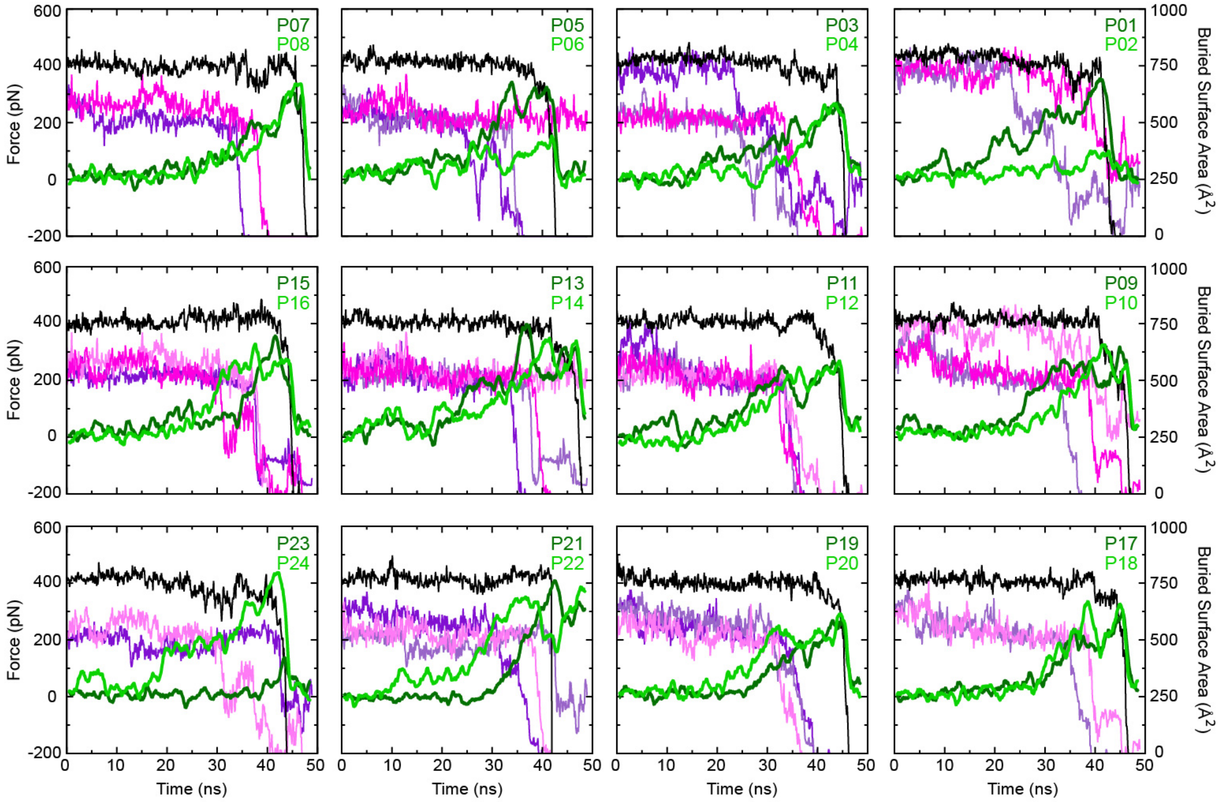


**FIGURE S5. Dynamics of *trans* and *cis* interactions during tensile stretching of the 24-CDH1 adherens junction.** Force versus time plots for constant-velocity stretching of individual CDH1 *trans* dimer pairs within the 24-CDH1 junction (0.5 nm/ns, simulation S1h) shown as in Fig. 2 *C*. Buried surface area as a function of time is overlaid (right scale) and shown for monomers involved in *trans* interactions (black), *cis* interaction on one side of the junction (pink), and *cis* interactions on the other side (purple). Brighter colors indicate the binding interaction is from EC1 and pale colors represent buried surface area coming from EC2.

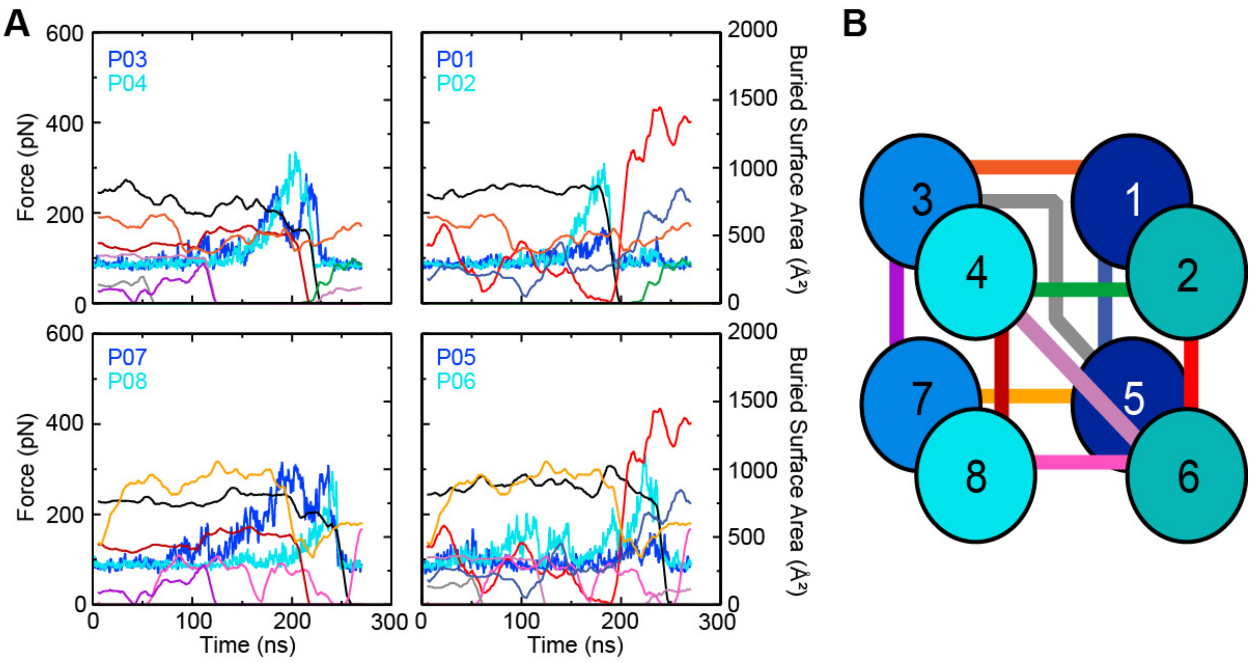


**FIGURE S6. Dynamics of *trans* and *cis* interactions during tensile stretching of the DSG2-DSC1 polarized desmosomal junction.** (*A*) Force versus time plots for constant-velocity stretching of individual DSG2-DSC1 *trans* dimers within the DSG2-DSC1 polarized junction (0.1 nm/ns simulation S3d) shown as in Fig. 4 *C*. Buried surface area as a function of time is overlaid (right scale) and shown for monomers involved in *trans* (black) and in *cis* (colored) interactions. There was no interaction between P01-P07 and P02-P08. (*B*) Schematic of the DSG2-DSC1 polarized lattice as shown in Fig. 4 *D* with the *cis*-interaction between monomers colored to match the buried surface area versus time plots in (*A*).

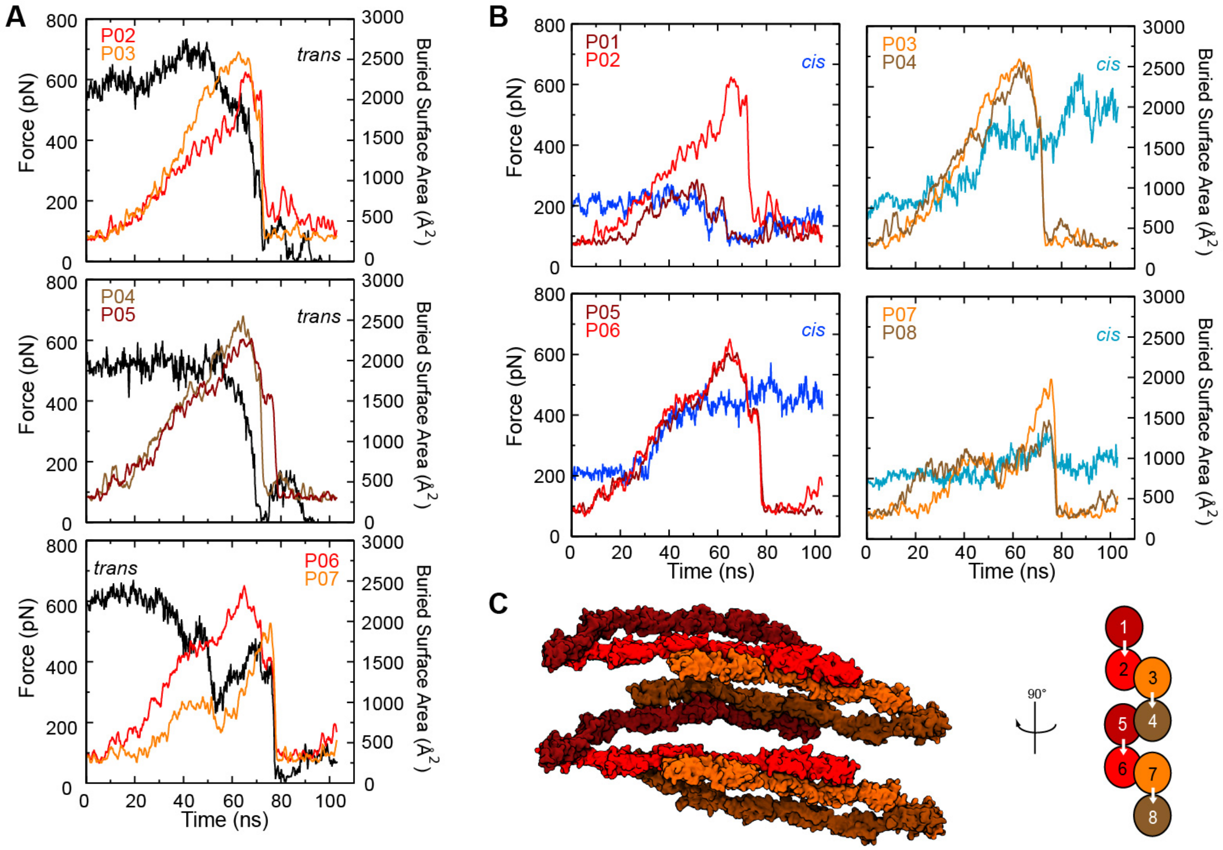


**FIGURE S7. Dynamics of *trans* and *cis* interactions during tensile stretching of the PCDHγB4 junction.** (*A*) Force versus time plots for constant velocity stretching of individual PCDHγB4 *trans* dimer pairs within the junction (0.1 nm/ns, simulation S5d). Buried surface area as a function of time is overlaid (right scale) and shown for dimers involved in *trans* interactions (black). Plots are arranged to reflect the position of PCDHγB4 *trans* dimers within the junction, as labeled in (*C*) (P01 – P08). (*B*) Force versus time plots for constant velocity stretching of individual PCDHγB4 *trans* dimer pairs grouped by *cis*-interacting pairs. Buried surface area as a function of time is overlaid (right scale) and shown for *cis* interactions (blues). Data labeled as in (*A*) and (*C*). (*C*) Schematic of the PCDHγB4 junction as seen in Fig. 7 *G*.
